# *Toxoplasma gondii* infection alters the recognition response of dendritic cells to stiffness substrate

**DOI:** 10.1101/2023.03.31.535021

**Authors:** ZhePeng Sun, Jing Liu, ZiFu Zhu, Zhu Ying, ZiHui Zhou, Qun Liu

**Affiliations:** National Animal Protozoa Laboratory, College of Veterinary Medicine, China Agricultural University, Beijing, 100083, China; Key Laboratory of Animal Epidemiology of the Ministry of Agriculture, College of Veterinary Medicine, China Agricultural University, Beijing, 100083, China

**Author notes:** Correspondence ZS JL ZZ ZY ZZ.

**Keywords:** **Keyword:** *Toxoplasma gondii*, dendritic cells, durotaxis, FAK, blood-brain-barrier

## Abstract

*Toxoplasma gondii* (*T.gondii*) hijacks host immune cells as ‘Trojan Horse’, and the infected cells accelerated the parasites dissemination. During acute infection, *T.gondii* specificity crosses the blood-brain-barrier to enter the brain. This selective mode of parasite transmission may be associated with the directed migration of infected immune cells. Immune cells follow various environmental cues for directional migration. However, the effect of *T.gondii* infection on the recognition of mechanical cues by immune cells remains unknown. Here, we examined the adhesion and migration of *T.gondii-*infected dendritic cells (DCs) on high and low stiffness substrates. We found that *T.gondii* infection alters the durotaxis migration of DCs. Infected DC exhibited stronger adhesion and lower migration on low stiffness substrates. In contrast to uninfected DCs, infected DCs migrated towards the low stiffness environment. TgWIP and TgROP17 co-regulate the F-actin structure of DCs and are involved in the formation of abnormal F-actin filaments. Rearrangement of the F-actin structure resulting from *T.gondii* infection regulates DCs’ abnormal recognition response to the mechanical cues. Recognition of DCs to the mechanical signals is independent of β2- integrin expression. Meanwhile, challenging DCs with *T.gondii* increased the phosphorylation of focal adhesion kinase (FAK). Treatment with a FAK inhibitor (VS- 6063) influences the recognition response of infected DCs. FAK inhibition in adoptively transferred infected DCs effectively prevents the dissemination of *T.gondii* to the brain. The data reveal that *T.gondii* infection inversely affects the durotaxis of DCs by altering the phosphorylation level of FAK and remodeling of F-actin structure. *T.gondii* utilizes the change in DCs’ durotaxis migration to accelerate the parasites crossing the blood-brain-barrier.

**Author Summary:** Immune cells travel through blood vessels and lymph vessels to various tissues, and respond to different types of environmental cues. Cells sense the cues and transmit these information to the cytoskeletal which induce directed cell migration towards or away from these signals. *T.gondii* infection remodeling the cytoskeletal of DCs which may cause abnormalities in these cues transduction. We found that *T.gondii* infection induces the formation of abnormal F-actin filaments in DCs, TgWIP and TgROP17 co-regulate the DCs’ F-actin structure. *T.gondii* infection increased the phosphorylation of FAK in DCs and has no effect with DCs surface β2-integrin expression. These reasons lead to alter the original durotaxis migration of DCs, and makes infected-DCs tend to stay in the low stiffness environment. Meanwhile, the recognition response of infected DC to mechanical signal determines the parasite rapid crossing the blood-brain-barrier.

## Introduction

*Toxoplasma gondii* (*T.gondii*) as an obligate intracellular parasite can infect the nucleated cells of almost all warm-blooded animals[1]. It is one of the few pathogens that have the ability to invade the central nervous system (CNS) by crossing the blood-brain-barrier (BBB)[2]. *T.gondii* hijacks host immune cells as Trojan horses and rapid disseminate to distant organs[3]. *T.gondii* drives immune cells to evade the host immune response and take nutrient from host cells [4]. During the acute stage of parasite infection, *T.gondii* exhibits specific organizational preferences, especially in the brain. Eventually, co-existing with the host by form bradyzoites in muscle and brain [5], that basis of chronic infection.

Immune cells respond to various types of environmental cues (chemotaxis, haptotaxis, durotaxis, topotaxis and galvanotaxis) which induce directed cell migration towards or away from the signals[6]. Through the regulation of these cues, immune cells quickly cross the vascular endothelial cells and reach to the specific tissues. The accelerated dissemination of *T.gondii* depends on the hijacking of immune cells. Recent studies have shown that *T.gondii* infection induces a hypermigratory phenotype of dendritic cells (DCs)[7], and infected DCs remain recognized chemical signal generated by CCL19[8]. It suggests that infected DCs still sensitive to environmental cues. Furthermore, the migration of infected DCs in vivo remains directional. Adoptive transfer of infected monocytes and DCs accelerates *T.gondii* dissemination to the CNS in mice[9]. The directed movement of infected monocytes or DCs to the brain may be associated with environmental cues. Interestingly, the brain is a softer tissue that provides a low stiffness microenvironment. This leads us to hypothesize that *T.gondii* infection affects immune cells to recognize environmental signals, especially mechanical cues (tissue stiffness).

Durotaxis refers to the tendency of cells to migrate toward regions where the substrate stiffness increases[10]. The execution of durotaxis depends on mechanosensing and – transduction which require the interactions between cell adhesion molecules and F-actin[11]. Integrins are important cell adhesion molecules, and mediate endothelial adherence which is the first step of transendothelial migration (TEM). Then, a fraction of F-actin is linked to the adopter molecules and binds dynamically to the specific ligands. *T.gondii* infection disrupts the F-actin structure of DC and dissolves the podosomes[12]. The remodeling of host F-actin is associated with parasite secreted proteins, such as TgWIP and TgROP17[13,14]. It indicated that *T.gondii* infection alters the interaction between adhesion molecules and F-actin which may shift the transduction of mechanical cues in infected cells. In addition, infected DCs that loss podosomes may require the focal adhesion or other adhesion molecules. Focal adhesion kinase (FAK) is a key component of focal adhesion complexes involved in cell-cell or cell-matrix adhesion, and regulates the signaling pathways initiated by integrins. *T.gondii* infection affects the phosphorylation level of FAK in DCs and reduces the expression of DC surface β1-integrin[15].

In recent years, research in the field has focused on the host cytoskeleton regulated by *T.gondii* infection. Yet, the recognition response of environmental cues by *T.gondii* infected cells remains unsolved. Here, we assessed the ability of infected DCs to recognize different stiffness substrates, and identify the role of infected DCs’ FAK in *T.gondii* dissemination in mice. Finally, we provided evidence for the effect of *T.gondii* infection on the durotaxis migration of DCs. The model of *T.gondii* infection also provides a new idea for the study of cell durotaxis migration.

## Results

### *T.gondii* infection leads to abnormal recognition of stiffness substrate

We used collagen I-coated polyacrylamide hydrogels of varying stiffness (elastic modulus ranging from 1 to 40 kPa by increasing the concentration of bis-acrylamide) to model different tissue stiffness[16]. *T.gondii* (RH-mCherry)-infected or uninfected primary murine bone marrow-derived DC (BMDC) were allowed to recognize onto different stiffness substrates (Figure 1a). The adhesion and migration of DC on stiffness substrate were indicators for cell mechanosensing and –transduction. Therefore, we compared the adhesion and migration tendency of infected or uninfected DCs in different stiffness substrates. Compared to the low stiffness substrate, uninfected DCs decreased migration distance (Figure 1b, c) and decelerated the average velocity (Figure 1d) of DC in response to high stiffness substrate. In addition, uninfected DCs exhibited stronger adhesion (Figure 1e) on high stiff. Strikingly, *T.gondii* (type I and II) infection inhibited adhesion (Figure 1e and S1d) and enhanced migration (Figure 1b-d and S1a-c) of DC on high stiffness substrate. As the substrate stiffness gradient decreases (the concentration of bis-acrylamide ranging from 0.26% to 0.03%), the adhesion of uninfected DCs gradually decreases while the infected DCs increase (Figure S1e). Collectively, these results suggest that *T.gondii* infection alters the durotaxis response of BMDC. Compared with uninfected DCs, infected DCs tend to stay in low stiffness.

**Figure 1.**
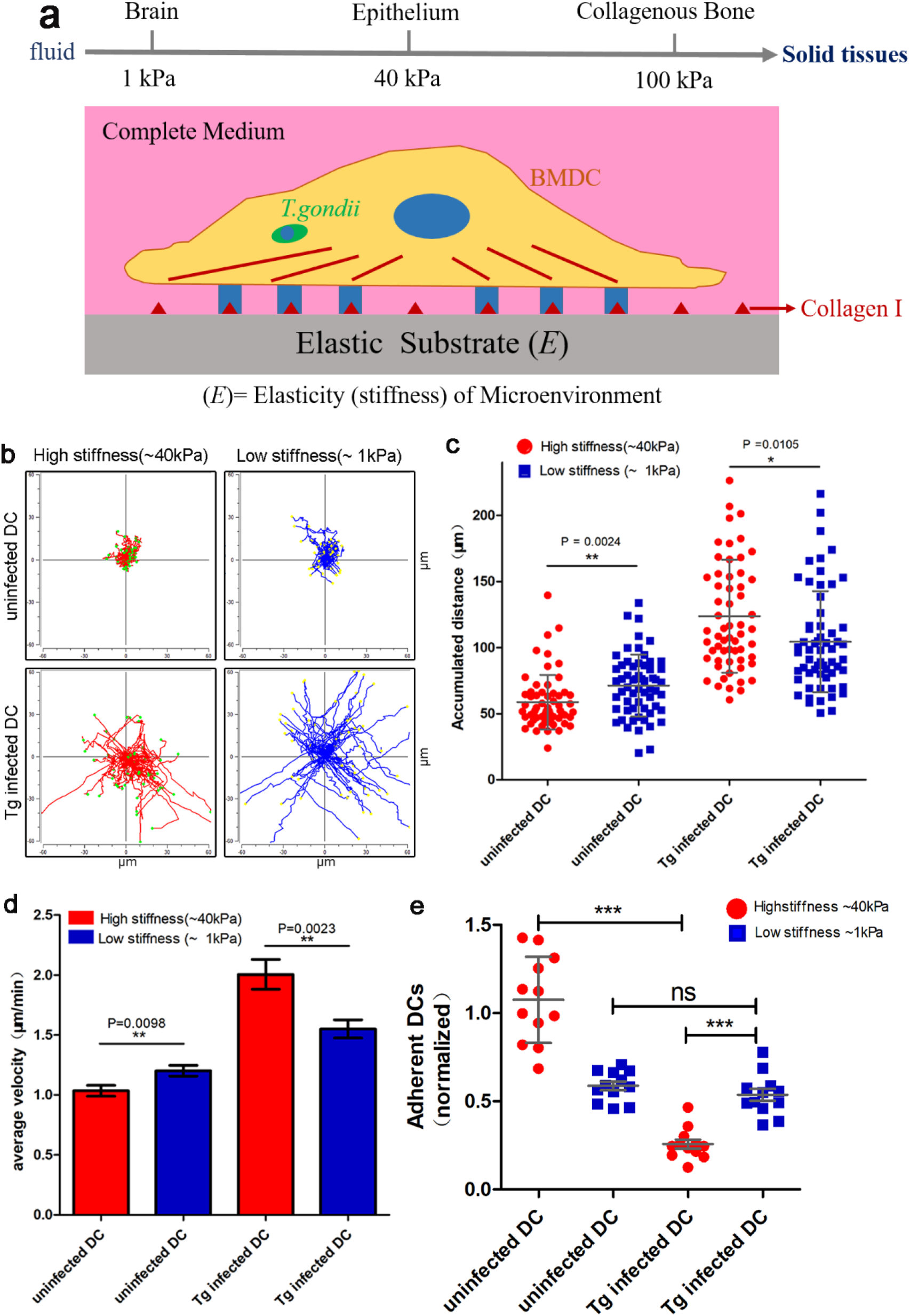
Recognition response of *T.gondii*-infected BMDCs to stiffness gradients substrate. (a) Schematic representation of migration and adhesion of uninfected or *T.gondii*-infected DCs on stiffness gradients substrate. The arrow on the top indicate the Elasticity (e) of different tissue microenvironment. (b) Mouse bone marrow-derived DCs were infected with RH*Δku80*-mCherry or left uninfected, the recognition of DCs on stiffness gradients was detected respectively as detailed in Materials and Methods. The migration of DCs were represented by motility plots and statistical analysis was performed for 60 cells. Data are representative of 3 independent experiments. (c) Accumulated migrated distances of uninfected or *T.gondii*-infected DCs on different stiffness substrate. The dot plot show the cell cumulative distance from 60 cells/group. (d) Cell velocity analyses of uninfected or *T.gondii*-infected DCs on different stiffness substrate. Graph show average velocity pooled from 3 independent experiments (n=3). (e) Cell adhesion analysis of uninfected or *T.gondii*-infected DCs on different stiffness substrate. Data were collected from three independent experiments. For all analyses, asterisks (*/**/***) indicate significant difference (\**p* < 0.05, ** *p* < 0.01, *** *p* < 0.001), ns (no significant difference). Normally distributed data were repeated measured with one-way ANOVA, Tukey’s test for multiple comparisons (c, e), and plotted as mean ± SD (d).

### *T.gondii* infection remodeling DC actin structure in both high and low stiffness substrate

*T.gondii* infection alters cell F-actin cytoskeleton which is exhibited as loss of podosome. This change in the F-actin cytoskeleton is the dynamic basis for the DC- hypermigratory phenotype[17]. However, in low stiffness environment, the F-actin structure of infected DCs remains unclear. To determine the actin structure of DC, we placed pre-infected BMDC to the high and low stiffness substrate for 6 h. The morphological results of F-actin showed that *T.gondii* (type I and II) infection causes podosome dissolution in both high and low stiff (Figure 2a and S2b). With loss of podosomes structure, *T.gondii* infected BMDC also appeared abnormal F-actin filaments. Different from traditional F-actin stress fibers traction by podosomes or focal adhesion (Figure S2a), the abnormal F-actin filaments without connected to the traction point (Figure 2a and S2b). In both high and low stiff, ∼70% uninfected BMDC contained podosomes while only ∼20% *T.gondii* (type I and II)-infected BMDC had podosomes (Figure 2b and S2c). The frequency of F-actin stress fibers was counted (for statistical purposes, we consider the abnormal F-actin filaments as ‘F-actin stress fibers’) by immunofluorescence, and only ∼20% uninfected BMDC appeared F-actin stress fibers. In comparison, ∼70% *T.gondii* (type I and II)-infected BMDC appeared F- actin stress fibers (Figure 2c and S2d). *T.gondii* infection also increased the cell area of BMDC in both high and low stiff. Similar to the uninfected BMDCs, infected DCs have a large size in high stiff compared to the low stiff (Figure 2d). Taken together, these results show that *T.gondii* infection leads to significant actin structure rearrangement in both high and low stiffness, manifested as loss of podosomes and the formation of abnormal F-actin stress fibers.

**Figure 2.**
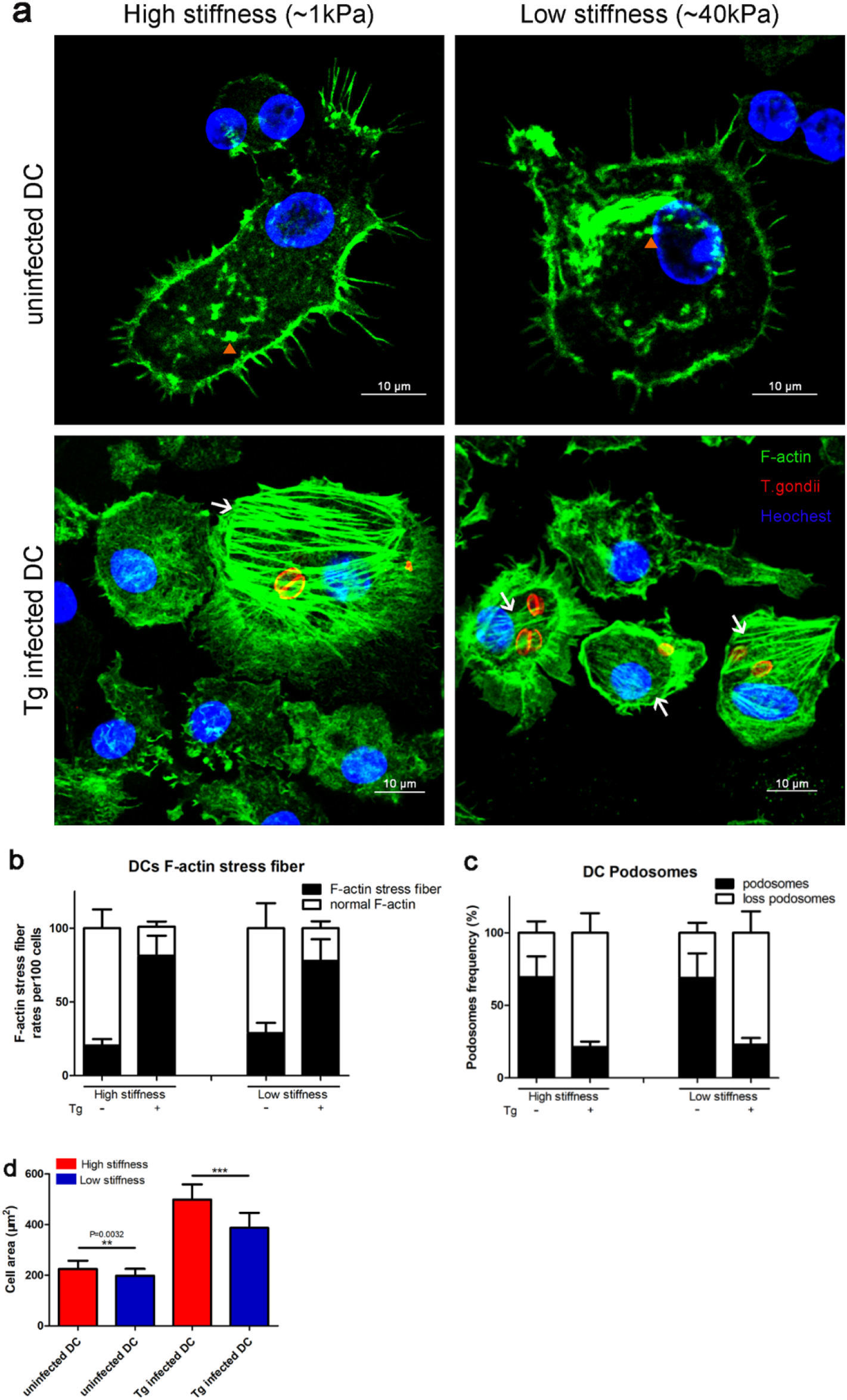
Morphological differences between uninfected DCs and *T.gondii*- infected DCs in stiffness gradients substrate. (a) DCs were uninfected or infected with RH*Δku80* for 6 h and then recognized on the stiffness gradients substrate for another 6 h. The cell podosomes (orange triangle) and F-actin stress fibers (white arrow) were stained with 488 Alexa Flour Phalloidin, the parasites were stained with anti-TgGAP45, and the nucleus were stained with Hoechest. Scale bars, 10 μm. (B and C) Quantification of the percentage of DCs appearing podosomes (b) and F-actin stress fibers (c) on the different stiffness substrate. Counted 100 cells for each group. (d) Cell area analyses of uninfected or *T.gondii*-infected DCs on the stiffness gradients substrate. The average cell area plotted as mean ± SD (n=3, count 100 cells per replicate). For all analyses, asterisks (*) indicate significant difference, repeated measured with one-way ANOVA and Turkey’s test for multiple comparisons.

### TgWIP and TgROP17 co-modulates parasitized DC actin structure

Previous studies have shown TgWIP modulates the aggregation of host F-actin structure by its WIRS/SH3 motifs[13]. The parasite activated host Rho-Rock signal pathway by secreting TgROP17. Amoeba-like movements of infected macrophages is associated with TgROP17 secretion[14]. These suggest that TgWIP and TgROP17 are key effectors that regulate host cell F-actin. The rearrangement of infected BMDC’s F- actin in both high and low stiff let us hypothesize that the formation of abnormal F- actin stress fibers is associated with TgWIP and TgROP17.

We used CRISPR/Cas9 to knock out the *wip* and *rop17* genes in RH*Δku80* and RH- mCherry. Successfully constructed single gene deletion strain *Δwip* and *Δrop17* (*Δwip-* mCherry, *Δrop17-*mCherry), and double-genes deletion strain *ΔwipΔrop17* and *ΔwipΔrop17-*mCherry (Figure S3a-b). The results of invasion assays, replication assays and plaque formation showed that there was no significant between gene deletion strains and parental strains (Figure 3f-g and S3c-e). Then we infected BMDC with parental, *Δwip*, *Δrop17*, and *ΔwipΔrop17* parasites for 3 h or12 h, and we found the formation of abnormal F-actin stress fibers is time-dependent to infection. Consistent with previous reports, infection of the *Δwip* parasite cannot induce the dissolution of host podosomes. As the *Δwip* parasite persistence in BMDC, abnormal F-actin stress fibers is unobservable. The infection of the *Δrop17* parasite leads BMDC to lose podosomes, but the abnormal F-actin stress fibers unable to formation. The infection of *ΔwipΔrop17* parasites has the same morphology with *Δwip* parasites infection (Figure 3a-e). To determine the formation of abnormal F-actin stress fibers, we infected BMDC with parental, *Δwip*, and *Δrop17* parasites, respectively. TgWIP and TgROP17 are rhoptries proteins, can secrete into the host cytoplasm with invasion of the parasite. Then, reinfection with parental, *Δrop17* and *Δwip* parasites after the first infection for 30 min (Figure S3f). The statistical results suggested that only the presence of both TgWIP and TgROP17 in BMDC could induce the production of abnormal F-actin stress fibers (Figure S3f-h). In summary, the host cytoskeleton regulates by *T.gondii* infection is time-dependent. DCs’ actin rearrangement of infected BMDC co-modulate by TgWIP and TgROP17. The appearance of abnormal F-actin stress fibers is associated to TgROP17, and the formation of F-actin stress fibers based on the dissolution of host podosomes.

**Figure 3.**
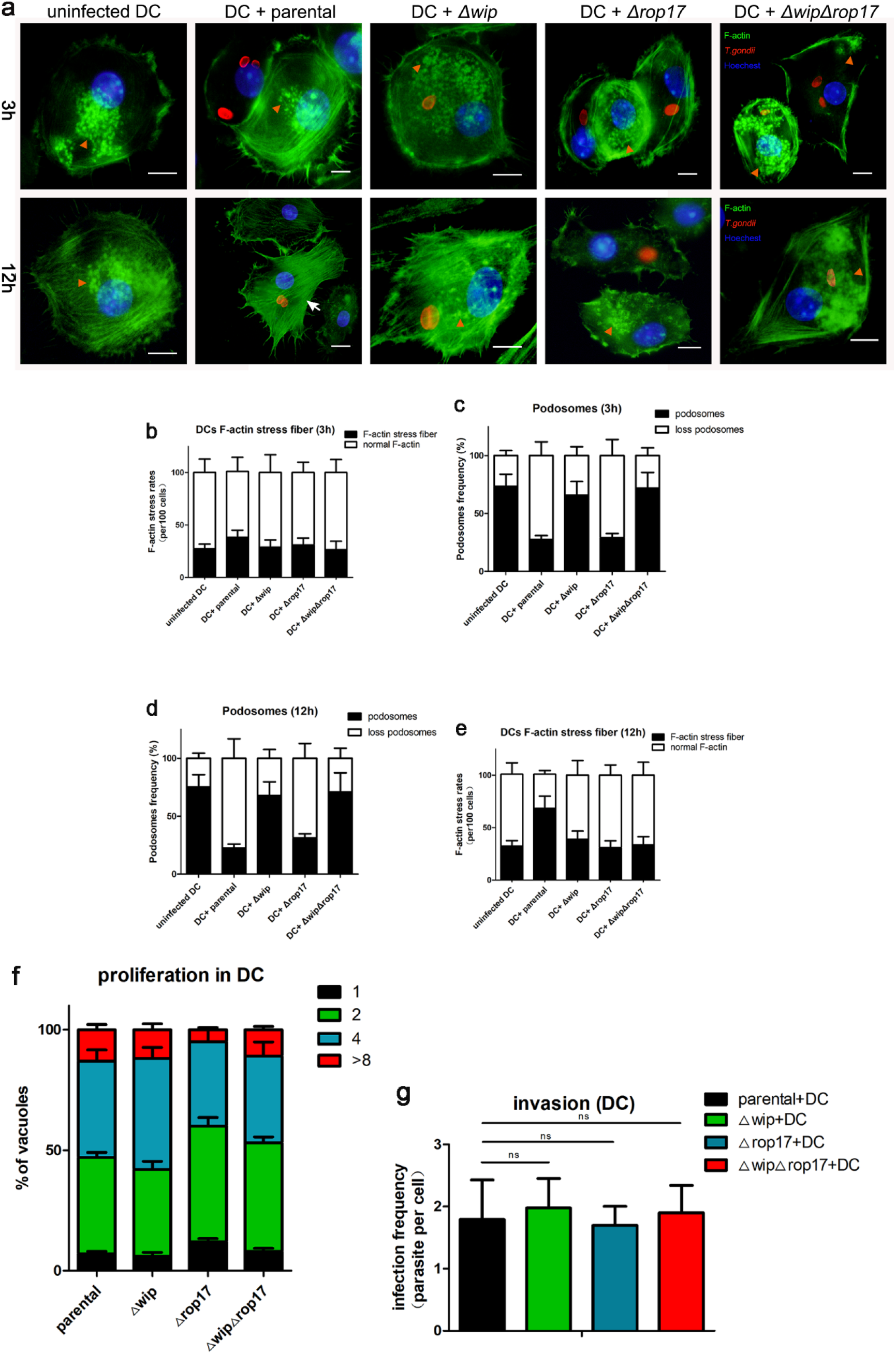
TgWIP and TgROP17 co-modulates DC actin structure. (a) DCs were respectively infected with RH*Δku80* (parental), *Δwip*, *Δrop17 or ΔwipΔrop17* parasites for 3 h or 12 h. DC’s F-actin structure were visualized by 488 Alexa Flour Phalloidin, the parasites were stained with anti-TgGAP45, and the nucleus were stained with Hoechest. Scale bars represent for 10 μm. (b and c) Quantification of the percentage of DCs containing podosomes on 3 h (b) or 12 h (c) pre-infected with RH*Δku80* (parental), *Δwip*, *Δrop17 or ΔwipΔrop17* parasites. Counted 100 cells for each group. (d and e) Quantification of the percentage of DCs appearing F-actin stress fibers on 3 h (d) or 12 h (e) pre-infected with parental, *Δwip*, *Δrop17 or ΔwipΔrop17* parasites. Counted 100 cells for each group. (f) The invasion assay of parental, *Δwip*, *Δrop17 or ΔwipΔrop17* parasites in DCs. Parasites infection frequency were counted by a total of 25 visual fields per group. Data are performed as mean ± SD (n=3), ns (no significant difference), one-way ANOVA, Dunnett’s multiple comparisons test. (g) The replication assay of parental, *Δwip*, *Δrop17 or ΔwipΔrop17* parasites in DCs. The number of parasites per vacuole was measured in DCs and a total of 100 vacuoles were analyzed per group. Data are performed as mean ± SD (n=3), ns (no significant difference) one-way ANOVA, Dunnett’s multiple comparisons test.

### *T.gondii* infection affects murine DC recognition to stiffness substrate by modulating mechanotransduction

The response of cells to mechanical cues (i.e, substrate stiffness) provided by the microenvironment is termed mechanotransduction[18]. Mechanotransduction requires mechanosensing, which is attributed to various cell surface molecules, such as integrin or cadherins[19,20]. The mechanical cues are then transmitted through the cell cytoskeleton. Infected DCs tend to stay in low stiff substrates, leading us to hypothesize that *T.gondii* infection may affect the cell mechanosensing and mechanotransduction. To determine cell mechanosensing in infected BMDC, we detected the expression of DC surface β2-integrin by flow cytometry. The expression of CD11a, CD11b and CD11c on BMDC cell surface has no significant between parental infected DCs and gene deletion (*Δwip*, *Δrop17* and *ΔwipΔrop17*) strains infected DCs (Figure 4a). We next examined the recognition of high and low stiff substrates by gene-deletion parasites-infected DCs. Similar to uninfected DCs, *Δwip* and *ΔwipΔrop17* parasites infected BMDC tend to rest on high stiff. In addition, *Δwip*, *Δrop17* and *ΔwipΔrop17* parasites infected BMDC did not exhibit the hypermigratory phenotype on both high and low stiffness substrate (Figure 4b-d). Compared to the parental parasite infection, the adhesion ability of *Δwip*, *Δrop17*, and *ΔwipΔrop17* parasites-infected BMDC increased on high stiff substrates (Figure 4e). However, in low stiffness environment, *T.gondii* infection has no significant effect on the adhesion ability of BMDC (Figure 4e). These results suggest that remodeling of the host cytoskeletal affect the recognition response of infected DC to the stiffness substrate. And this recognition response is independent with β2-integrin expression.

**Figure 4.**
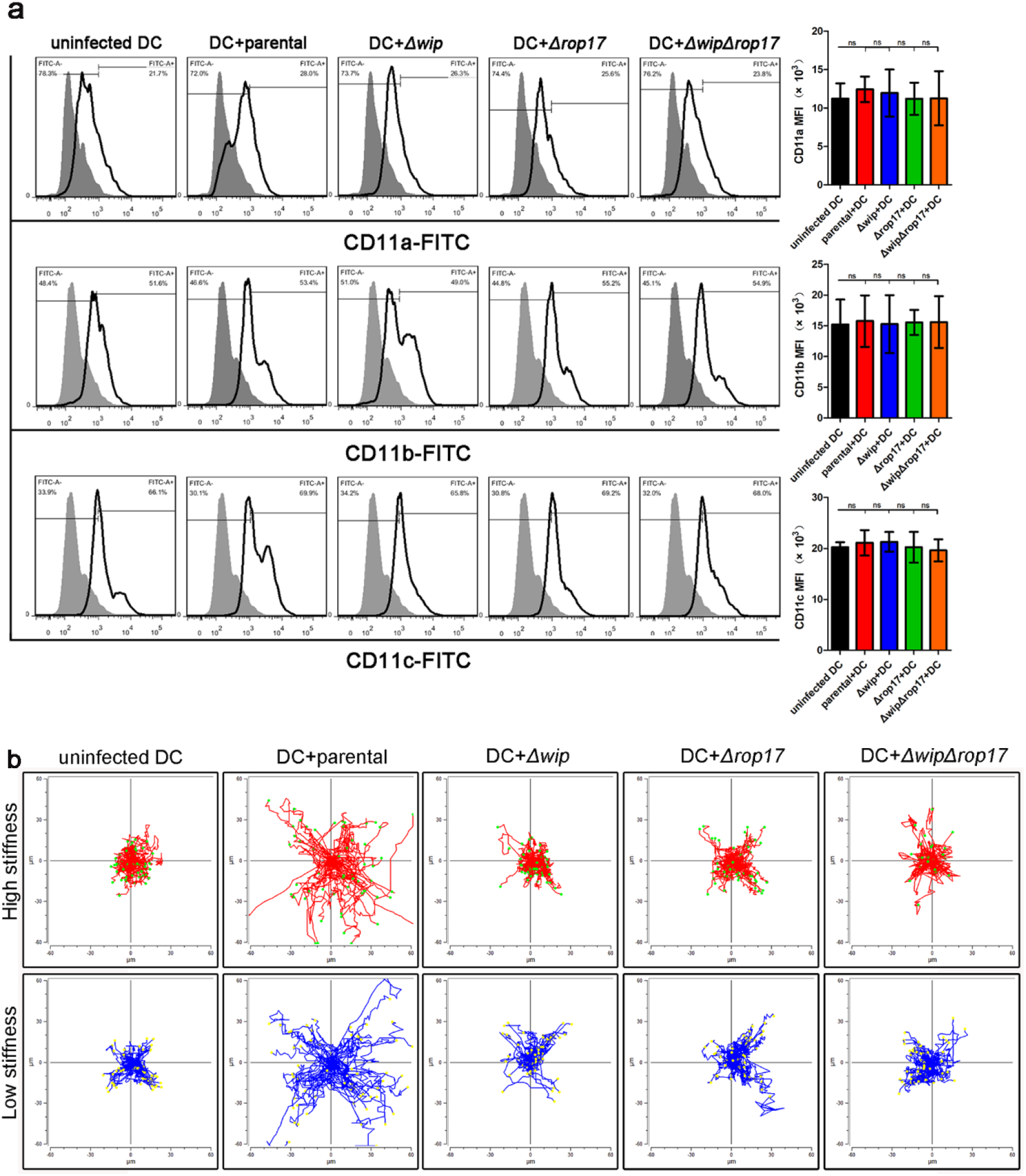
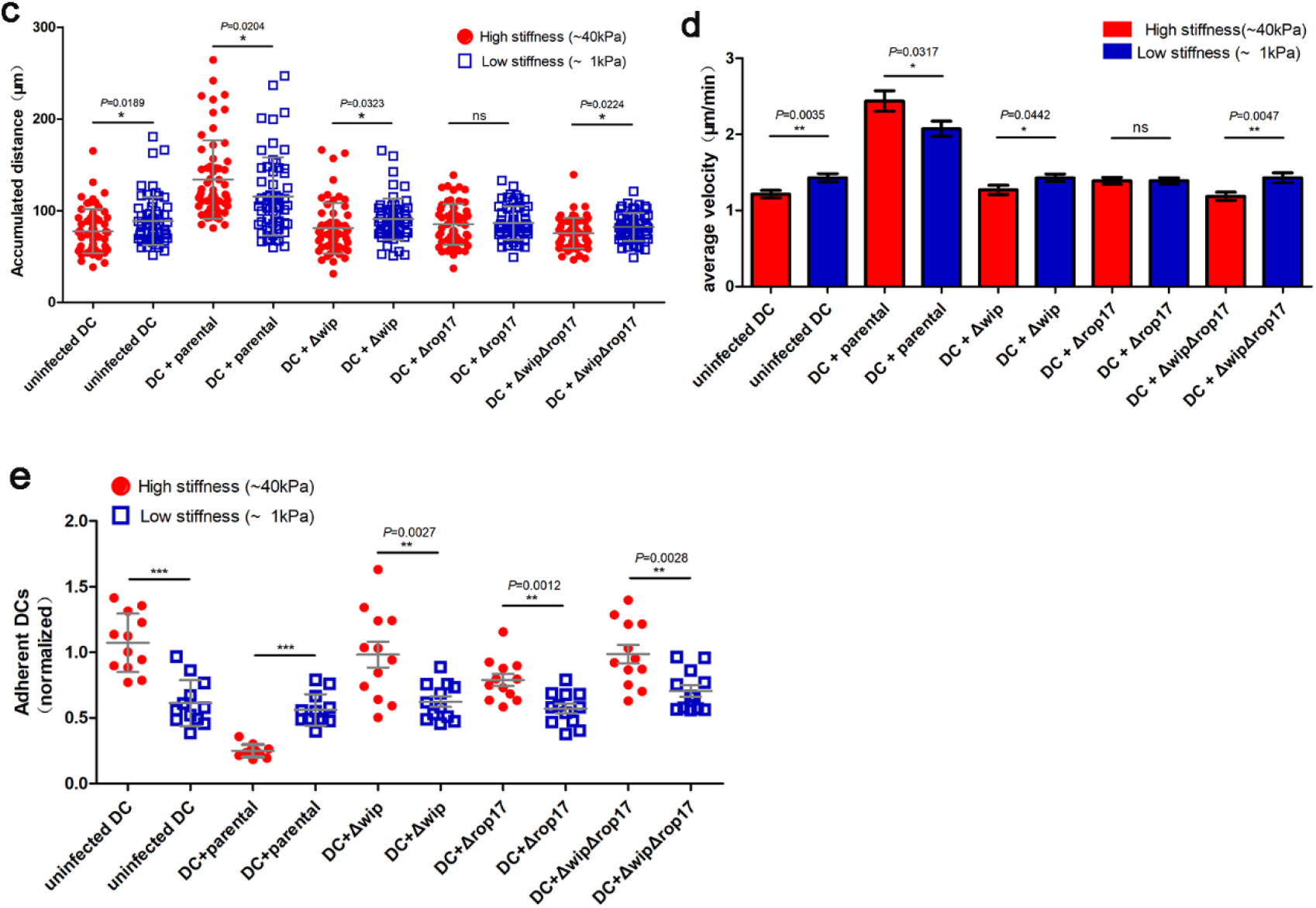
Effect of abnormal actin structure on DC recognition substrate stiffness. (a) Flow cytometric analysis of the expression of uninfected or *T.gondii* (RH-mcherry, *Δwip-*mcherry, *Δrop17-*mcherry *or ΔwipΔrop17-*mcherry parasites)-infected DCs surface β2-integrin (CD11a, CD11b, CD11c). Histograms show gates used to define CD11a^+^, CD11b^+^ or CD11c^+^ cells (right). Graph show the median fluorescence intensity (MFI) of CD11a, CD11b or CD11c (n=3 independent experiments). (b) DCs were respectively infected with RH*Δku80*-mcherry (parental), *Δwip-*mcherry, *Δrop17-* mcherry *or ΔwipΔrop17-*mcherry parasites for 6 h and then recognized on the stiffness gradients substrate. Migration of infected-DCs were analyzed by motility plots. Data were pooled from three independent experiments. (c) Accumulated migrated distances of uninfected or *T.gondii*-infected DCs on different stiffness substrate. The dot plot show the cell cumulative distance from 60 cells/group. Asterisks (*) indicate significant difference, with one-way ANOVA and Turkey’s test for multiple comparisons. (d) Cell velocity analyses of uninfected or *T.gondii*-infected DCs on different stiffness substrate. Average velocity were performed as mean ± SD (n=3), asterisks (*) indicate significant difference: one-way ANOVA and Dunnett’s multiple comparisons test. (e) Cell adhesion analyses of uninfected or *T.gondii*-infected DCs on different stiffness substrate. Data were collected from three independent experiments and were analyzed with one-way ANOVA and Dunnett’s multiple comparisons test.

### FAK is involved in the recognition response of infected DCs to stiffness substrates

The cells form unstable adhesion in low stiffness environment, which keeps the cells in a low adhesion state. Unstable adhesion is regulated by a variety of adhesion molecules, including integrin-related adhesion [21]. Different from the high stiffness substrate, the adhesion ability of infected DCs (loss podosomes) in low stiffness substrate does not have a significant reduction (Figure 1e). Therefore, we suspect that *T.gondii* infection may affect the function of other adhesion molecules. We used inhibitors to screen out adhesion-related molecules associated with cell mechanotransduction. The results show that FAK-inhibitor (defactinib, VS-6063) significantly reduces the adhesion of infected DCs in low stiffness (Figure S4a). *T.gondii* infection increases the phosphorylation level of FAK (T397) in BMDC, and the phosphorylation level of DCs’ FAK is independent with host F-actin structure rearrangement (Figure 5a). Next, we detected the adhesion of *Δwip*, *Δrop17*, and *ΔwipΔrop17* parasites-infected DCs treated with FAK-inhibitor. Compared to uninfected DCs, the treatment of VS-6063 significantly reduce the adhesion of parental and *Δrop17* parasites-infected DCs in low stiffness substrates (Figure 5b). In addition, in high stiffness environment, the FAK-inhibitor lessens the adhesion of infected or uninfected DCs (Figure S4b). Moreover, treatment with the FAK inhibitor decreased migration distance and decelerated the average velocity in both high and low stiffness substrates (Figure 5c-e and S4c-e). It suggests the phosphorylation of FAK resists the effect of podosomes dissolution on adhesion in the low stiff environment. Furthermore, the abnormal recognition response of infected DCs was partially restored by FAK-inhibitor treatment (Figure 5f-g). These results suggest that FAK is a critical effector for infected DCs to recognize the mechanical cues.

**Figure 5.**
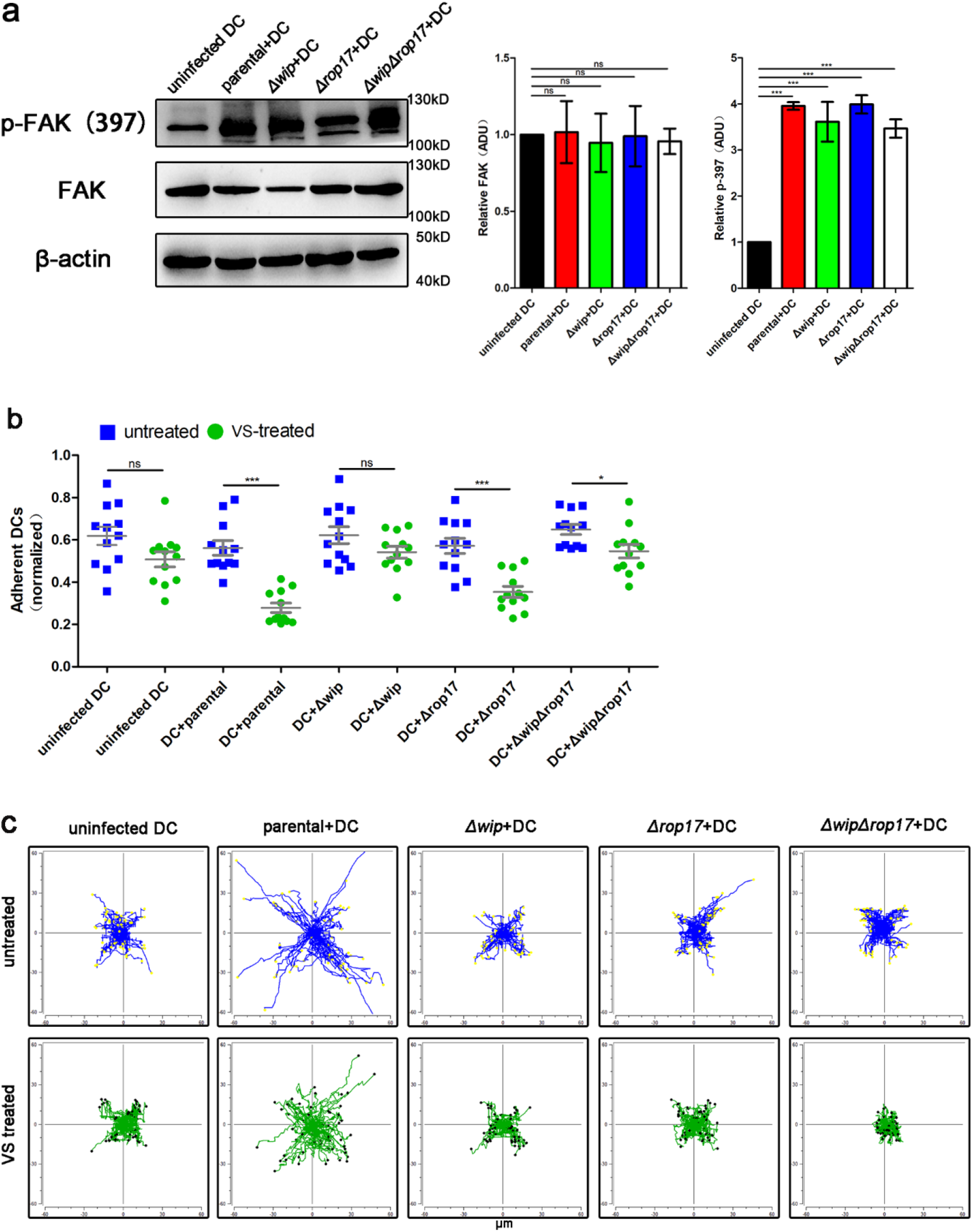
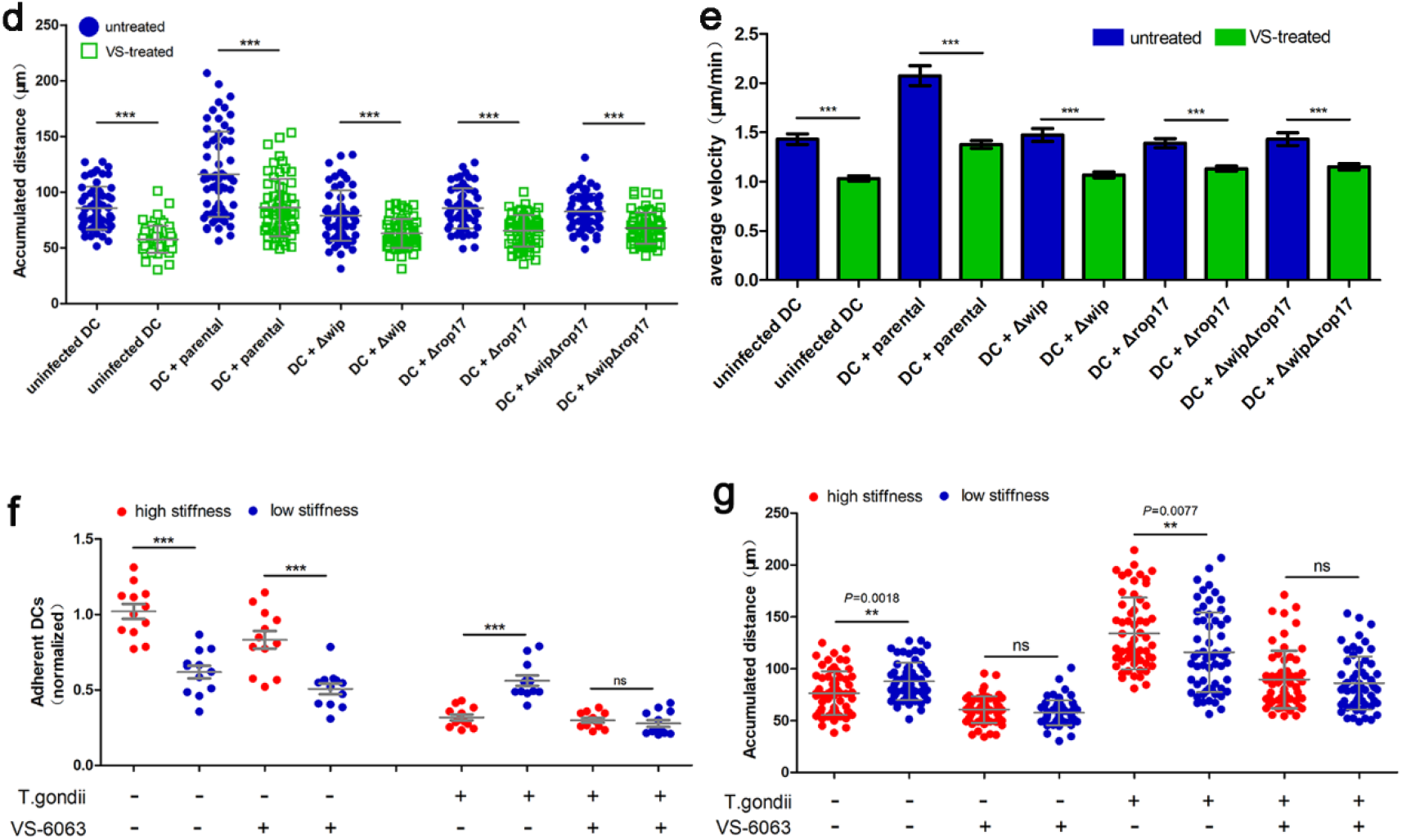
FAK regulate the DCs recognition response to the low stiffness substrate. (a) Western blot analysis of total FAK protein and phosphorylated FAK (T397) expression in *T.gondii*-infected DCs. Graph shows FAK/p-397 normalized to β-actin expression. Arbitary denitometry unit (ADU) were analyzed from three independent biological replicates. (b) Cell adhesion analyses of VS-6063 (FAK-inhibitor) treated with *T.gondii* (parental, *Δwip*, *Δrop17 or ΔwipΔrop17* parasites)-infected DCs on low stiffness substrate. DCs were pre-treated or untreated with VS-6063 (20 μM) for 6 h and then infected with parasites at MOI 3. After parasites infection, the cell adhesion of *T.gondii*-infected DCs were detected. Data were collected from three independent experiments (n=3) and were analyzed with one-way ANOVA and Dunnett’s multiple comparisons test. (c) DCs treated with VS and infected with parasites. Migration of DCs on low stiffness substrate were represented by motility plots. Data are representative of three independent experiments. Accumulated migrated distances (d) and cell velocity (e) analyses of VS pre-treated and untreated DCs on low stiffness substrate. Uninfected or *T.gondii*-infected DCs average velocity were peformed as mean ± SD (n=3), asterisks (*) indicate significant difference: one-way ANOVA, Dunnett’s multiple comparisons test. Cell adhesion (f) and cell migration (g) analyses of VS-6063 (FAK-inhibitor) treated with infected DCs or uninfected DCs in high and low stiffness. The tendency of DCs adhesion to the different stiffness substrates were detected. Data were collected from three independent experiments (n=3) and were analyzed with one-way ANOVA and Dunnett’s multiple comparisons test.

### FAK inhibition in adoptively transferred infected murine DC reduces the parasitic load and dissemination of *T.gondii* in vivo

Previous studies have demonstrated that adoptive transfer of *T.gondii*-infected DCs accelerates the dissemination and exacerbates parasite infection compared to fresh tachyzoites [22]. To assess the impacted of FAK inhibition on infected DCs in vivo dissemination. We determined the mouse virulence of *Δwip*, *Δrop17*, and *ΔwipΔrop17* parasites, and found that there are no significant difference between parental and gene deletion strains (Figure S5a). Subsequently, we inoculated i.p. with parasites (*Δwip*, *Δrop17*, and *ΔwipΔrop17*)-infected DCs and found the mouse virulence of infected DCs has no considerable difference between parental and gene deletion (Figure S5b). Next, we pre-treated DC with FAK-inhibitor and then infected with parental and *ΔwipΔrop17* parasite. However, treatment of FAK-inhibitor had no effect on the mouse survival curve (Figure 6a). Due to the excessive motility of the type I parasite, we then used the type II parasite (Pru) to examine the dissemination of infected DCs treated with FAK-inhibitor. Survival curve show VS treatment delayed the mouse death (Figure 6b). To determine the presence of parasites in different organs, we collected the spleen, brain, liver, and peritoneum fluid on the 7 days after infection and quantified the parasitic loads by qPCR. Overall, we found that challenged with infected DCs had higher parasitic loads in the spleen, brain, and liver tissue compared to VS pre-treated DC infected with the parasite. However, the parasitic loads of peritoneum fluid have no substantial difference between VS-treated and untreated DC (Figure 6c). Altogether, these results indicated that treatment of infected DCs with FAK-inhibitor results in a significant reduction in the parasitic load of target tissue and dissemination of *T.gondii* during the course of infection in mice.

**Figure 6.**
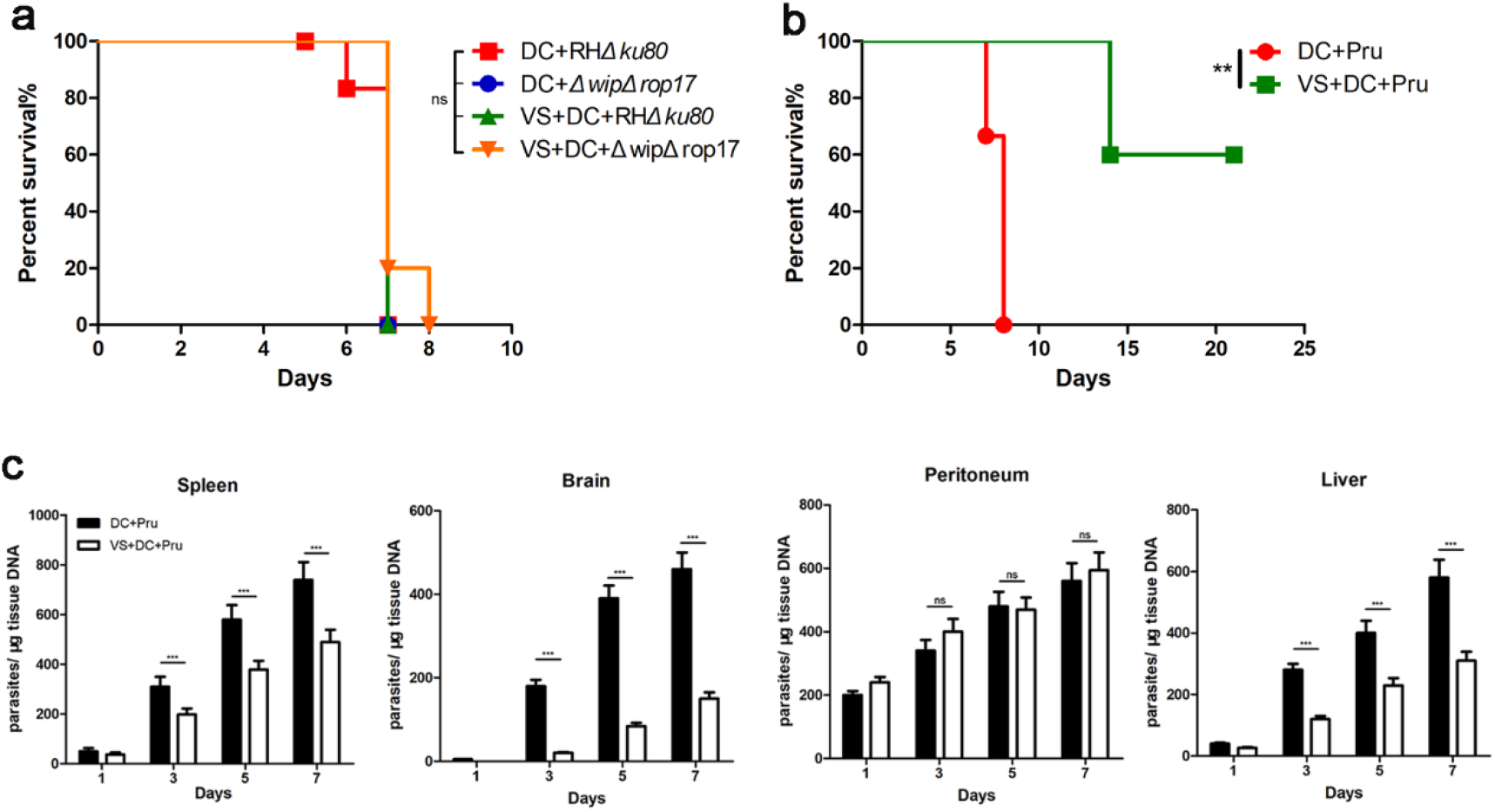
Adoptive transfer of *T.gondii*-infected DCs treated with FAK inhibitor decreased parasite load in vivo. (a) Female BALB/c mice were infected intraperitoneally with 200 parasites (RH*Δku80* or *ΔwipΔrop17* parasites) or 200 *T.gondii* (RH*Δku80* or *ΔwipΔrop17* parasites)-infected DC (n=2, 5 mice for each group). (b) Female BALB/c mice i.p. infected with 2,000 wild-type Pru-infected DCs or 2,000 Pru-infected DCs pre-treated with FAK-inhibitor. Asterisks (*) indicate significant difference, ns (no significant difference). Survival curve (a, b) were analyzed with log rank Mantel-Cox test, Gehan-Breslow-Wilcoxon test. (c) Parasitic load (Pru) in spleen, brain, peritoneum and liver on day 1-7 post inoculation quantified by qPCR as indicated under Materials and Methods. BALB/c mice were infected with 5×10^4^ of Pru-infected DCs or 5×10^4^ Pru-infected DCs pre-treated with FAK-inhibitor. Standard curves for tissue 28sRNA and Tg 529 were established. Tissue parasites burden was expressed as the content the number of parasites per microgram tissue DNA. Significant differences in parasite load between groups were observed in spleen, brain and liver (Kruskal-Wallis test). Non-significant difference were detected in peritoneum (Kruskal-Wallis test, P≥0.05).

### Treatment with FAK inhibitor reduces the frequency of infected DCs crossing the BBB

To determine the effects of FAK on the directional migration of infected DCs in mice. We isolated mouse brain microvessels on 3, 5, and 7 days after mouse infection. Counted the frequency of parasites’ appearance in brain microvessels (Figure 7a). 3 day after infection, *T.gondii* appeared in the brain microvessels of adoptively transferred Pru-infected DCs. Infected DCs pre-treated with FAK-inhibitor significant decrease the dissemination of *T.gondii* (Figure 7b). The ratio of intravascular and extravascular parasites show that infected DCs enhance TEM. Parasites located in the brain microvessels quickly across the BBB by untreated DC (Figure 7c). The distance of extravascular parasites localized to the nearest blood vessel has been counted, no significant difference between untreated and VS-treated DC infected with Pru (Figure 7d). In period infection of Pru-infected DCs, *T.gondii*-infected cells appear around the brain microvessel. However, in post-infection, *T.gondii* infected cells are invisible nearby the vessels (Figure 7e). In summary, infected DCs treatment with FAK-inhibitor reduces the efficiency of parasites across the BBB.

**Figure 7.**
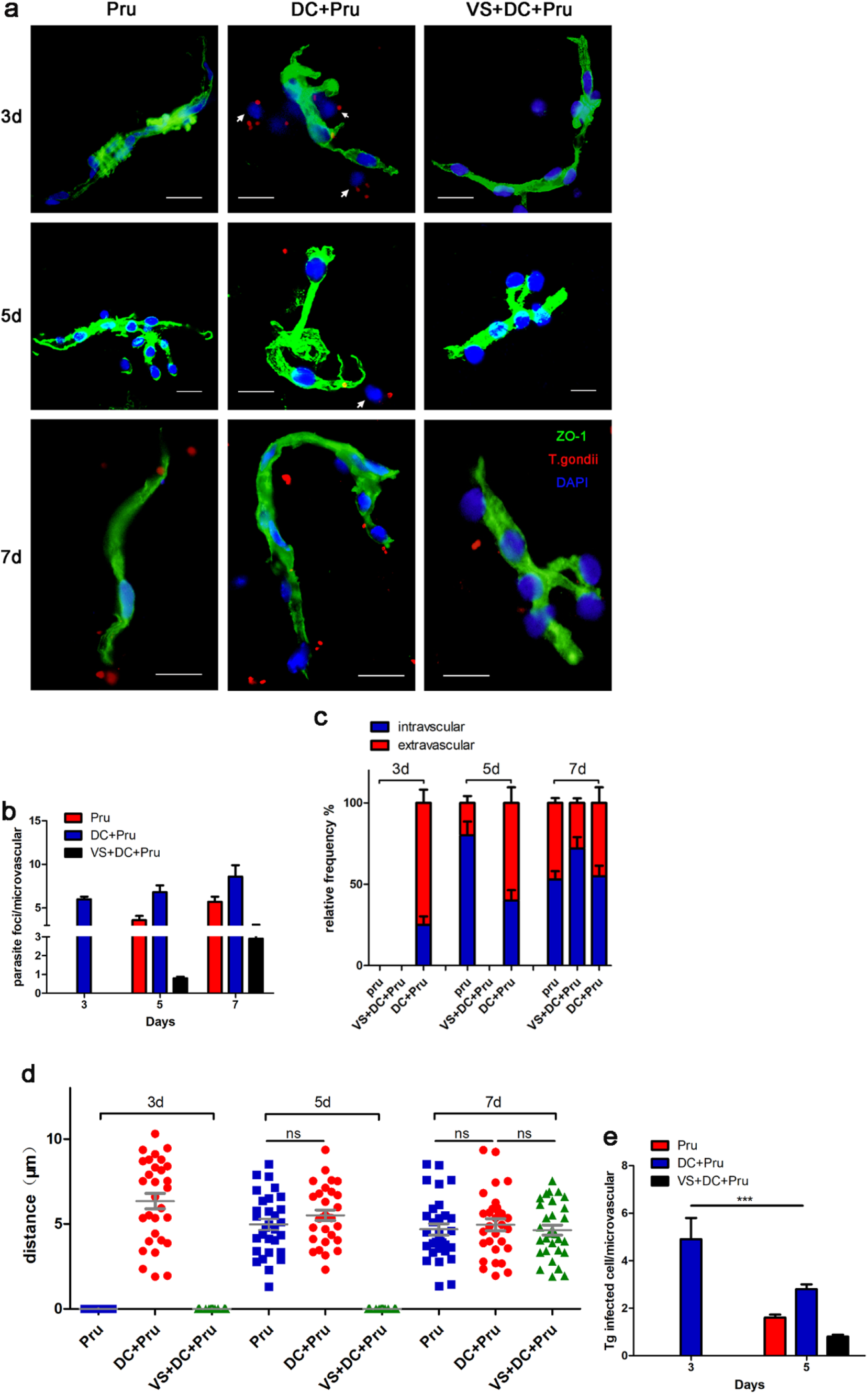
FAK inhibitor decreased *T.gondii*-infected DCs dissemination to Blood-brain barrier. (a) Female CD-1mice were infected with 5×10^4^ Pru, 5×10^4^ Pru-infected DCs or 5×10^4^ Pru-infected DCs pre-treated with VS-6063. Mouse brain microvessels were isolated on 3, 5, 7 day and identified by BBB marker anti-ZO-1. Nuclear stained with DAPI and Pru were stained with TgGAP45. Arrowhead indicates extravascular cell infected with *T.gondii*. Scale bars, 20μm. (b) The total numbers of parasite foci (TgGAP45-Cy3^+^) located extravascular or intravascular related to the numbers of brain microvessels on 3, 5, 7 day post-inoculation. Data were collected from 60 microvessels per group. (c) CD-1 mice were inoculated with Pru, Pru-infected DC or Pru-infected DC pre-treated with VS-6063. Relative frequency (%) of parasite foci related to microvessels localization at 3, 5, 7 day post-inoculation. (d) The distance of extravascular parasite to the nearest vascular branching. n=10 independent foci. ns, no significant difference, Student’s t-test. (e) The total numbers of *T.gondii*-infected cells appeared around the brain microvessels. Data were collected from 60 microvessels per group. Asterisks (*) indicate significant difference with two-way ANOVA, Turkey’s test for multiple comparisons.

## Discussion

In our study, we select collagen I-coated polyacrylamide hydrogels as stiffness substrate for cell recognition. Because the porosity of hydrogel is similar to the tissue microenvironment[23]. And compared to the substrate formed by lysyl oxidase (LOXL1-LOLX4) crosslink collagen[24], the hydrogel substrate has more comprehensive stiffness range[25]. Immune cells can respond to substrate stiffness which induces directed cell migration toward to the high stiffness substrate[26,27]. A recent study showed that human neutrophils has increased spreading and lower migration ability, but stronger adhesion ability on high stiffness substrate[28]. High local endothelial cell (EC) stiffness promotes leukocyte adhesion and benefits leukocyte for search the sites that permit TEM. In low local EC stiffness, the reverse of leukocytes spreading provide the condition for rapid TEM[29,30]. We found the migration toward in murine BMDC conforms to typical durotaxis. However, *T.gondii-* infected DCs show migration tendency to low stiffness microenvironment. These suggest that *T.gondii* infected DCs has an abnormal recognition response to the mechanical cues.

The change of cell durotaxis in *T.gondii-*infected DCs is associate with cell mechanosensing and mechanotransduction. Cell mechanosensing is achieved through various transmembrane adhesion molecules, including integrin or cadherins. The recognition of leukocytes to stiffness signal is regulated by these adhesion molecules[31]. Previous report has shown that *T.gondii* infection reduces the expression of monocytes and DCs β1-integrins[12]. However, the expression of infected DCs surface β2-integrins is still controversial. Our results show *T.gondii* infection has no effect on the expression of DC surface β2-integrins subunit CD11a, CD11b, and CD11c. It indicated that the change of DC durotaxis is not dependent on the β2-integrins recognition. In addition, whether the mechanosensing of infected DCs regulated by β1-integrins or cadherins still needs further study. Integrins are force-generating mechanoreceptors that derived force signals into the actin-skeleton[32]. The cytoplasmic tail of integrin tilts within the cell membrane as actomyosin contraction, resulting in the position of integrins shift on the cell membrane[33]. As cell movement progresses, the length and angle of inclination to integrins on the cell membrane are also important indicators of mechanosensing. However, in our trial, we failed to detect these data. The recent result from Lisa L. Drewry et.al show that *T.gondii* infection reduced the monocytes surface integrin contact area to substrate[14]. It suggests that *T.gondii* infection tends to inhibit integrin contact with the substrate rather than its expression.

After recognition of mechanical cues, further transmission of signals requires mechanotransduction which induces F-actin or myosin-based contraction[34,35]. Inhibition of F-actin dynamics in human neutrophils impairs cell spreading and reduces cell TEM[36]. The contraction of F-actin is the basis of leukocytes to response the substrate stiffness. However, *T.gondii* infection directly affects regular stiffness recognition by rearrangement of the host F-actin structure. TgWIP is associated with the degradation of podosomes, and our experiments also confirmed this phenotype. As the infection progressed, we found that abnormal F-actin filaments appeared in the infected DCs and were related to the secretion of TgROP17. This F-actin filament is similar to the F-actin stress fibers, but without an explicit tension point. The production of these excessive stress fibers dependent on the secretion of TgWIP and TgROP17. These stress fibers affects the normal mechanotransduction and generate traction force which induces infected DCs hypermotility. Furthermore, podosomes also participate in the secretion of matrix metalloproteinases (MMP)[37]. The matrix-degrading enzymes can soften the substrate and activate the downstream mechanical landscape[38]. *T.gondii* infection promotes DC secretion of TIMP-1 which acts as an inhibitor of MMP [15]. This encourages infected DCs to contact the stiffer microenvironment and sculpt the stiffer mechanical cues. Moreover, actin filaments connected with the nucleus by LINC (linker of nucleoskeleton and cytoskeleton) complex which acts as a sensor for mechanotransduction load on either side of the nucleus[39]. Whether the abnormal actin filaments in infected DC link to the LINC complex is still unknown. And more work will be required to detect these ideas. The adhesion and migration of *Δwip*, *Δrop17* parasites*-*infected DCs demonstrate that parasite regulates durotaxis recognition of DCs through the cytoskeletal remodeling. There are also many parasite secrete effectors that regulate the migration of immune cells, such as Tg14-3-3, TgGRA28, and TgMYR1[40-42]. The regulation between parasite effectors and cell durotaxis needs more research.

*T.gondii* infection results in the dissolution of DC podosomes which associate with cell adhesion. We report that loss podosomes induce lower adhesion of DC in high stiffness substrate. But in low stiffness environment, podosome dissolution has no effect on DC adhesion. It may be related to the construction of unstable focal adhesion by cells in low stiffness substrate. Focal adhesions is the first step in TEM, and the partial components of podosome structure are involved in the formation of focal adhesions[43]. Not all cells form stable focal adhesion, such as monocytes, but they still sensitivity to the substrate stiffness[44]. *T.gondii* infection may activate the cytoplasmic region of adhesion-related molecules or downstream molecules. Strikingly, we found FAK is a critical molecule that determines the adhesion of infected DCs in the low stiff environment. FAK is a component of focal adhesion that recruits by extracellular matrix ligands and regulates the cell-cell or cell-matrix adhesion. The auto-phosphorylates of FAK occurs during the cell adhesion and migration[45]. The deficiency of FAK leads to increased permeability of EC and unstable tight junction between ECs[46]. Meanwhile, the phosphorylation level of FAK is critical for cell to transmit mechanical cues. Our data reveals that *T.gondii* infection increase the level of FAK phosphorylation in DC which is independent F-actin rearrangement. We pre-treated FAK inhibitor with infected DCs, and found FAK inhibitors can effectively reduce the adhesion of infected DC in low stiffness. Increased level of FAK phosphorylation in *Δwip*, *Δrop17* parasites*-* infected DC demonstrate that excessive phosphorylation of FAK was not associated with F-actin rearrangement in DC. Treatment of FAK inhibitor prevents the sensing of mechanical cues from infected DC and partially reverses the infected DCs’ durotaxis. Our findings suggest that adoptively transferred infected DCs increase the frequency of parasites appearing in cerebral microvessels. After *T.gondii* infection, DC exhibits amoeboid-like migration termed hypermotility, and similar migration enhancement occurs in both macrophages and monocytes[47]. During TEM, EC stiffness provides a physical basis for leukocyte adhesion and migration. The sensitivity of leukocytes to stiffness microenvironment determines the efficiency of TEM. The abnormal recognition of *T.gondii*-infected DCs to stiffness may guide parasites to migrate into low stiffness tissue. FAK inhibitor pre-treated DC prevent parasites transmission to the brain, and also provide evidences for the influence of DC abnormal durotaxis for parasites’ quick dissemination. Furthermore, stiffness is only one factor determines the TEM. The adhesion receptor (such as P-selectin and ICAM1) on the surface of EC can capture leukocytes and enhance the adhesion of leukocytes, and is also essential for TEM[48]. FAK is a crucial factor in quick dissemination of *T.gondii* into deep tissue during the acute infection.

In conclusion, *T.gondii* infection reverses DC durotaxis by rearrangement of the F-actin structure. The abnormal durotaxis of *T.gondii*-infected DCs is crucial for parasite toward migration to the brain. The regulation of the cytoskeleton by *T.gondii* provide a new sight for cell durotaxis research. Meanwhile, this toward migration to low stiffness microenvironment of infected DCs also provide a new idea for *T.gondii* dissemination during acute infection.

## Supplementary information

Supplementary Figure 1 to 5

## Author contributions

QL and ZS conceived the study. ZS and JL performed the experiments. ZS, ZY, and ZZ analyzed the data. ZS designed and wrote the manuscript. ZZ helped in animal experiments. ZY and ZZ helped in manuscript writing. All authors read and approved the final manuscript.

## Ethical statement

The raising and handling of animals are in compliance with the recommendations in the Guide for the Care and Use of Laboratory Animals in China. All experiments were approved by the Institutional Animal Care and Use Committee of China Agricultural University (under the certificate of Beijing Laboratory Animal employee ID: 18049). The surviving mice were injected with atropine (0.02mg/kg) at 30dpi, and were humanely sacrificed by cervical dislocation.

## Acknowledgements

We are also grateful to Prof. Bang Shen (Huazhong Agricultural University, China) for providing the CRISPR/CAS9 vector. We are appreciation to Prof. Shaojun Long (China Agricultural University, China) for the support of confocal microscopy imaging.

## Inclusion and diversity

Data supporting the conclusions of this article are included within the article. We support diverse, equitable, and inclusive conduct of research.

## Materials and Methods

### Cell lines and parasite cultures

HFF (Human Foreskin fibroblast) monolayers and Vero (African Green Monkey Kidney Cell) cell lines were cultured with 10% and 8% fetal bovine serum (FBS, Gibco, USA) DMEM (Dulbecco’s Modified Eagle’s Medium) complete medium. Tachyzoites from RH (type I) and Pru (type II) were maintained by serial 3-day passage in HFF or serial 2-day passage in Vero.

### Animals

C57BL/6 mice (female, 5 weeks old), BALB/c mice (female, 4 to 5 weeks old) and CD- 1 mice (female, 4 to 5 weeks old) were purchased from Vital River Laboratory Animal Technology Co., Ltd (Beijing, China) and maintained under pathogen-free conditions with ad libitum access to clean food and water.

### Generation of mouse bone marrow-derived DC (BMDC)

Mouse bone marrow-derived DC were generated by cytokine induction [49]. In brief, femurs from 7 to 12 weeks old C57BL/6 mice were washed through with phosphate-buffered saline (PBS). Bone marrow was collected and cultured in RPMI-1640 complete medium (CM) containing 20ng/ml recombinant mouse GM-CSF (Peprotech). Subsequently, bone marrow were maintained in culture medium at 37 ℃, 5% CO_2_ with the addition of 20 ng/mL rm GM–CSF and 10 ng/mL rm IL-4 (Clone-cloud). The CM replaced every 2 days, by aspirating 1 mL of media and replacing with 1 mL of fresh CM supplemented with cytokines. Adherent cell were harvested on day 6 for subsequent experiment.

### Plasmid construction and parasite transfection

Plasmids were assembled from DNA fragment by DNA ligase (Vazyme). The details of all the primers used in this study are listed in Table 1.The CRISPR/Cas9 system was used to generate gene knockout strains and endogenous genomic locus strains. CRISPR/Cas9 plasmids used in this study were derived from the single-guide RNA (sgRNA) plasmid pSAG1:Cas9-GFP,U6:sgRNA:UPRT, preserved in the Key Laboratory of Animal Parasitology (Beijing, China).

Freshly harvested parasites were transfected with plasmid, using protocols previously described [50]. In brief, 1.5×10^7^ freshly tachyzoites were maintained in 350 μl cytomix buffer, mixed with 50 μl purified plasmid (CRISPR/Cas9 plasmid) and DNA fragment (amplification products containing homologous DNA fragments) in a 4-mm Gene Pulser and electrotransformed by Gene Pulser Xcell system (Bio-Rad). Transgenic parasites were selected with pyrimidine (1μM) or chloramphenicol (20μM). Monoclonal strains were isolated by dilution on HFF monolayers grown in 96-well plates.

### Immunofluorescence assay

HFFs/BMDCs were cultured on glass coverslips in 12-well plate. After parasite infection, 4% of paraformaldehyde was used to fix host cells for 30 minutes, subsequently permeabilized with 0.25% of Triton-X 100 for 20 minutes and blocked with 3% BSA for 30 minutes. Primary antibodies were incubated for 1 hour and washed with PBS. After secondary antibodies incubated for 1 hour, the nuclei of host cells and parasites were stained with Hoechst and host cell cytoskeleton were stained with Alexa Fluor^TM^ 488 Phalloidin (ThermoFisher, USA). Mouse anti-TgSAG1 and rabbit anti-TgGAP45 were stored in our laboratory.

Images were observed by the Olympus IX70 Inverted Microscope and captured by Olympus DP Controller software. Cytoskeleton were viewed using laser scanning confocal microscopy (Leica TCS SP8 X, Germany). Podosomes were identified and quantified as described in previous report[51], 100 cells were analyzed for each experiment. Mean fluorescence intensity (MFI) and cell area were analyzed by Fuji image (ImageJ, http://imagej-nih.gov/ij/).

### Preparation of polyacrylamide hydrogels substrate

Construction of stiffness gradients polyacrylamide hydrogels substrate were following a method described previously [16].In brief, prepared a large piece of coverglass (45mm×50mm) as a base plate and flamed in alcohol burner repeatedly. After cooling at room temperature, coverglass was soaked in 0.1M NaOH solution and air dried. Pipetted 150-200 μl of 3-aminopropyltrime thoxysilane (Sigma-Aldrich) on one side of the glass and spread evenly onto the surface. After 5 minutes, the coverglass was washed and soaked in distilled H_2_O. Then immersed the glass in a solution of 5% glutaraldehyde in PBS for 30 minutes and air dried. Stiffness gradients polyacrylamide were mixed by acrylamide and bis-acrylamide, containing 10% acrylamide and bis-acrylamide concentrations ranging from 0.26 to 0.03%. Different concentrations of acrylamide/bis-acrylamide mixture (add sodium supersulphate and TEMED) was then placed on the coverglass respectively, and covered with a small circular piece of coverglass (∼15mm diameter) as roof. After gel polymerization, the small coverglass was removed and rinsed the gel with 200 mM Hepes (pH 8.5). The gel was incubated with 50mM heterobifunctional crosslinkers Sulfo-SANPAH (Sigma-Aldrich) in 200 mM Hepes, and exposed to the UV light for photoactivation. After 5 minutes photoactivation, the Sulfo-SANPAH removed from the gel, and washed with200 mM Hepes. Contain 0.2 mg/ml rat tail collagen I (Sigma-Aldrich) solution was then layered onto the stiffness gradients polyacrylamide substrate and reacted overnight at 4℃. Before cell culture, the gel was soaked in the CM for 40 minutes at 37℃.

### Cell migration and cell adhesion assay

For cell migration assay, polyacrylamide gel was placed in a 24-well plate coated with poly-L-lysine (Sigma-Aldrich) respectively. After the gel was fully fitted to the cell plate, CM was added and incubated at 37 ℃ for 30 min. DCs were infected with *T.gondii* (MOI of 3 with Rh-mCherry,*Δwip*-mCherry,*Δrop17*-mCherry or*ΔwipΔrop17*- mCherry) for 6 h. In each well, 5×10^5^ uninfeted-DCs or infected-DCs were mixed with rat collagen I (1 mg/ml) and placed on the stiffness gradients substrate for 15-30 minutes. Confirm the cells were contact with the gel under the microscope. Cell trajectories were measured for 1h, 1frame/min at 10×magnification by Z1 Observer (Zeiss). Cell migration was analyzed by Manual Tracking (ImageJ plugin), count 60 cells per experimental group and time-lapse images were incorporated into stakes. Cell average velocities and accumulated distance were analyzed by Chemotaxis and migration tool v.2.0 (Ibidi). Infected cells were defined by mCherry co-localization.

For cell adhesion assay, 5×10^4^ uninfeted-DCs or infected-DCs per well were placed in a 96-well plate pre-treated with polyacrylamide gel. DC were allowed to adhere on the stiffness gradients polyacrylamide substrate for 45 min at 37℃. Then, non-adherent DCs were vigorously rinsed by plate washer (Bio-rad). The adherent was measured by Cell-Couning-Kit-8 (CCK-8, MCE). Samples were detected by multifunctional microplate reader (biotek synergy h1), and Gen5 software (Biotek) was used to quantify the number of adherent cells. The experiments were repeated three separate times with 6 technical replicates for each condition.

### Invasion assay and replication assay

For invasion assay, the DCs/HFFs spread into 24-well plate and added 1×10^6^ freshly tachyzoite in each well [52]. Placed the cell plate in cell incubator for 30 minutes. Washed the cell plate 3 times by PBS, and fixed with 4% paraformaldehyde. Non-invaded parasites were stained with anti-rabbit SAG1 before permeabilization, and the invaded parasites were detected with anti-mouse IMC1 after permeabilization. Goat-anti rabbit IgG (H+L)-FITC and goat-anti mouse IgG (H+L)-Cy3 were used for secondary antibodies, Hoechst was used to stain the nucleus. Parasites labelled in green were scored as the extracellular, and marked with red were counted as intracellular. Independent invasion assays were performed three times, and each trial quantified 25 fields to evaluate the invasion efficiency. The invasion ratio was calculated by invaded parasites/total parasites (invaded + non-invaded).

For the replication assay, a total of 1×10^5^ freshly tachyzoites were added into DC or HFF cells in 12-well plate. After incubated in cell incubator for 30 minutes, cells were washed 3 times by PBS. Fresh medium was replaced into the cell plate and incubated for another 22 hours. Anti-rabbit GAP45 and hoechest were used for IFA and a total of 100 parasite vacuoles were counted under fluorescence microscopy for the replication assay. The experiments were repeated three separate times.

### Flow cytometry

DC surface integrin was analyzed by flow cytometry. The cells were pre-infected with T.gondii for 6h, then collected and diluted to 5×10^5^ cells/100 μL with PBS. Each sample was measured in triplicate. FITC-labeled anti-CD11a, anti-CD11b, anti-CD11c (Abcam) were added into the cell suspension to a final concentration of 5μg/mL, and incubated for 20 min at 25℃ in the dark. Mouse IgG2a isotype was used for control antibodies. The cells were washed with PBS for twice. Then tested all cell samples by flow cytometry (FACSCalibur, BD Bioscience). The result analyzed by Flowjo 7.6 software (flowjo software, Inc, LLL, USA).

### Western blot

To quantify the protein levels of infected-DCs FAK and phosphorylated FAK (T397), DCs were infected with RH*Δku80*, *Δwip*, *Δrop17*, *ΔwipΔrop17* parasites for 6h. *T.gondii* infected-DCs were collected and lysed in RIPA buffer (50mM Tris, 150mM NaCl, 0.1% Triton, 0.1% SDS and 0.5% deoxycholic acid) with cocktail protease and phosphatase inhibitors (Roche) at 4℃ condition. Proteins were separated using 7.5% SDS-PAGE gels and blotted onto a PVDF membrane (Millipore, 0.45μm) and blocked followed by incubation with anti-FAK (CST), anti-β-actin (CST) and anti-phosphorylation-FAK T397 (CST). After primary antibodies were incubated for 1 hour and washed with TBS, secondary antibodies (anti-rabbit IgG HRP) incubated for another 1 hour. Proteins were detected by enhanced chemiluminescence (Thermo Fisher) in a BioRad ChemiDoc MP. Densitometry analyses was performed using Fuji image (ImageJ, NIH, USA).

### Adoptive transfers of DC

Before *T.gondii infection*, DC were treated with VS (FAK-phosphorylation inhibitor, 20μM) for 6h. Then, DC were challenged with freshly egressed tachyzoites for 6 h (MOI: 1 at I type parasite strain and 2 at II type parasite strain), and non-invaded parasites were washed by PBS. Following the cells were collected and diluted in PBS, 5×10^4^ cells were adoptively transferred into male BALB/c or CD-1 mice. Before injection, inhibitor groups were treated with an additional 20μM VS for 1h. The cells were injected i.p. into BALB/c or CD-1 mice.

Six day post adoptively transformation CD-1 mice were euthanized, and collected peritoneal fluid and organs. Parasite quantitative detection in mice was performed by qPCR. In brief, prepared tissue slurry and DNA was extracted by DNA Extraction kit (Aide Lai Biotechnology Co. Ltd., Beijing, China). Primers were designed to quantitatively detect mouse 28S rRNA and *T.gondii* 529 gene by qPCR. Standard curves for 28sRNA and Tg529 were established. Mouse survival was detected by BALB/c mice. Five mice per group were observed daily with ad-libitum access to clean food and water and recorded the number of mouse deaths.

### Isolation of murine brain microvessels

Cerebral microvessels were isolated from CD-1mice as described (). In brief, mouse brain were gently separated with conjunctiva forceps and minced with scalpel-blade. The brain fragments were maintained in D-hanks (Sigma-Aldrich) solution and then homogenized by passing through a 23-gauge needle. Brain homogenate was digested by 1mg/ml type VI collagenase (Gibco) at 37℃ for 1h. Then, tissue homogenate was mixed in an equal volume of 70% Percoll (Sigma-Aldrich) and centrifuged at 5,000g for 15min at 4℃. After removal the top layer (myelin layer), the pellet was resuspend in D-hanks and passed through a 75μm cell strainer. The cell strainer was then flushed with PBS to collect the mouse brain microvessels. The brain microvessels were identified by ZO-1 expression.

### Statistical analyses

Statistical analyses were performed using GraphPad Prism software (version 8.1.1). All data are presented as average±standard deviation (SD). For all the calculation *P*<0.05 are considered as significant. For two group comparisons, tow-tailed Student’s t-test was used for normal distribution. For more than three groups, data with a normal distribution were analyzed by one-way ANOVA, followed by Dunnett’s test.

---

**Supplementary figure 1.**
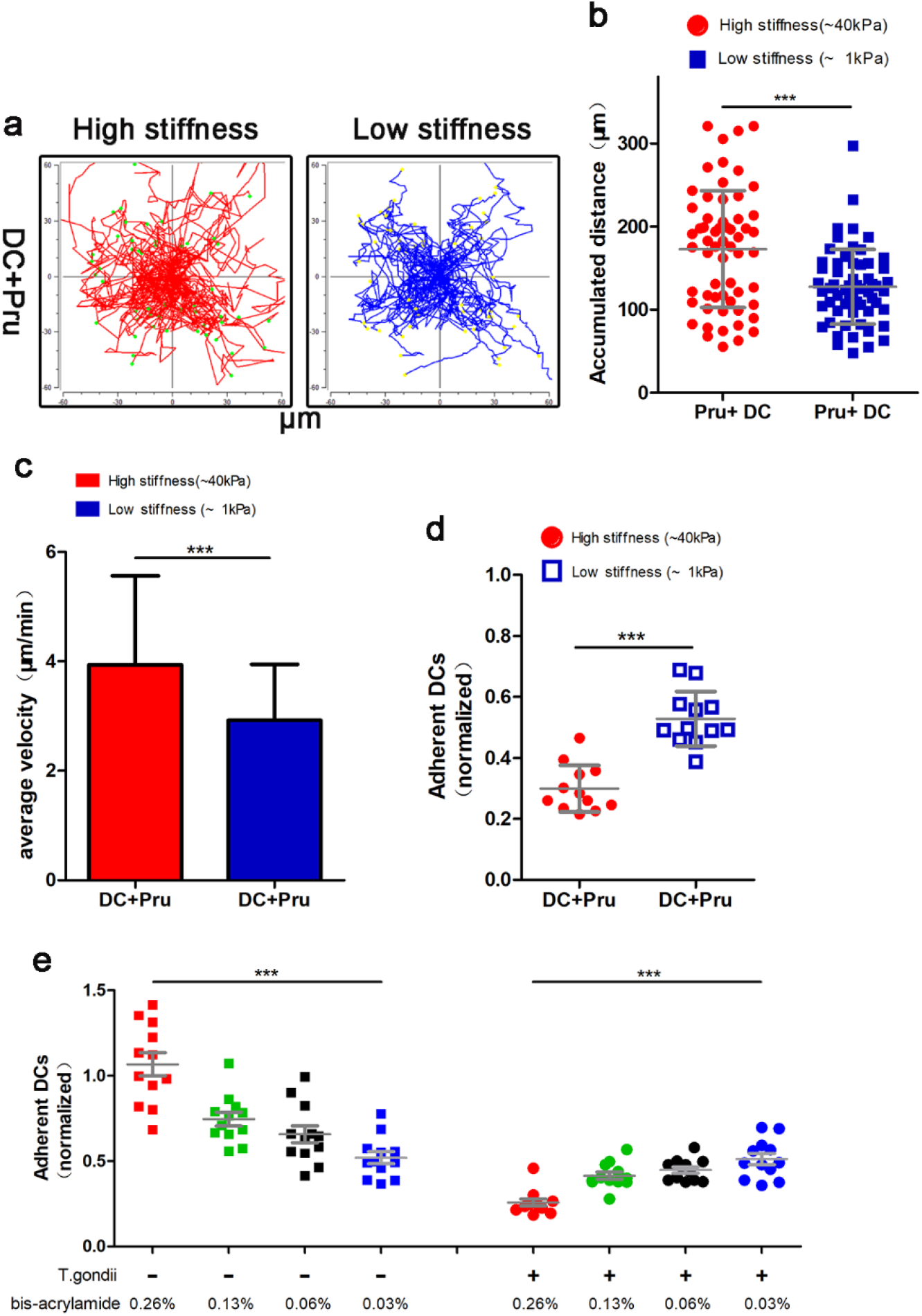
Recognition response of Pru-infected DCs to stiffness gradients substrate. BMDCs were infected with Pru, the recognition of DCs on stiffness gradients was detected respectively. The analysis of motility plots (a), accumulated migrated distances (b), cell average velocity (c) and cell adhesion (d) were collected from three independent experiments. (e) Cell adhesion analysis of uninfected or *T.gondii*-infected DCs on stiffness gradients substrate (the concentration gradient of bis-acrylamide is 0.26%, 013%, 0.06% and 0.03%).

**Supplementary figure 2.**
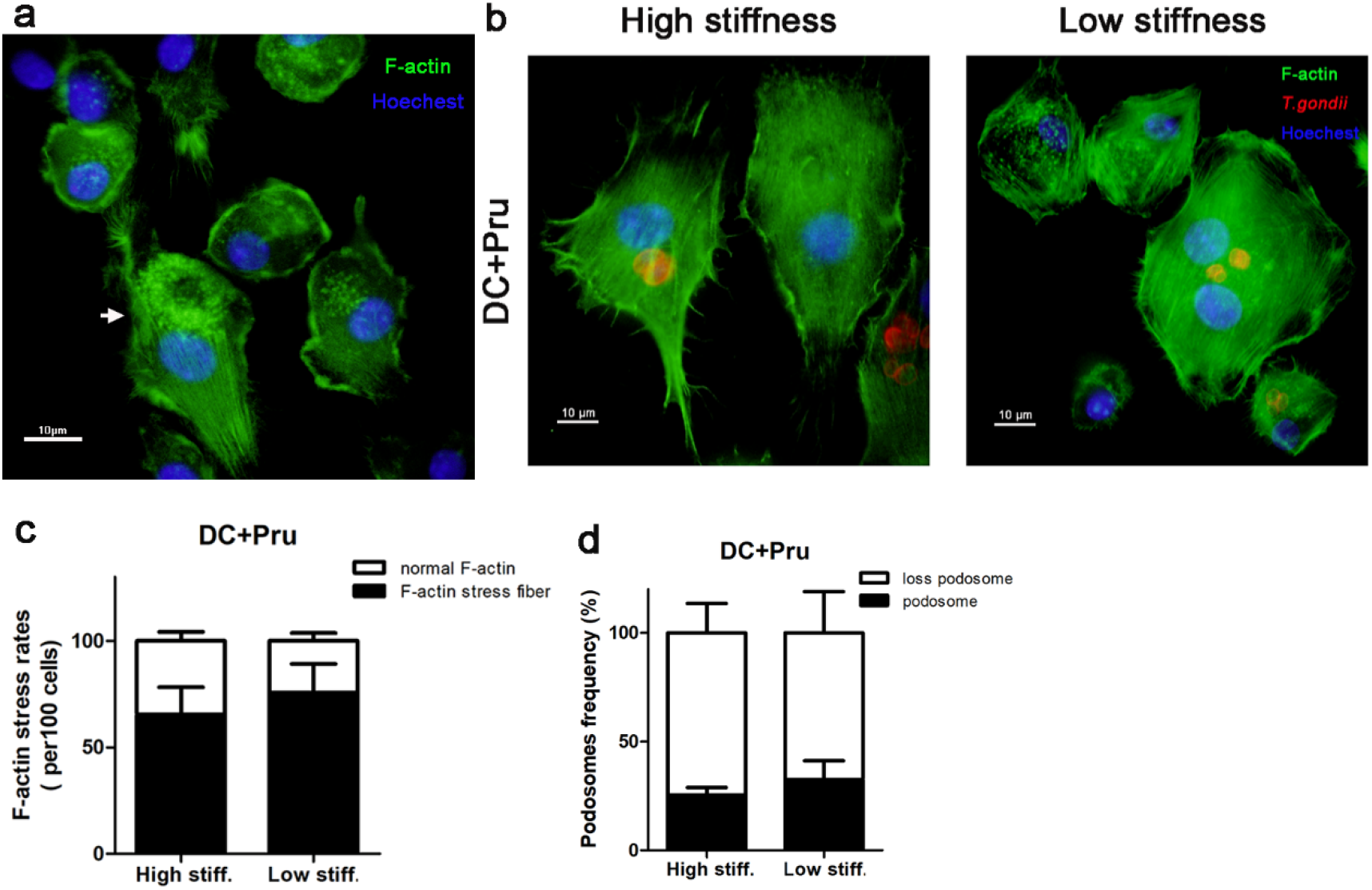
Actin structure of Pru-infected DC on stiffness gradients substrate. (a) The F-actin structure of uninfected DCs. F-actin stress were stained with 488 Alexa Flour Phalloidin, the nucleus were stained with Hoechest. White arrow show the combination of podosome structure and stress fibers. Morphological of Pru-infected DC actin structure (b, c and d) were analyzed with one-way ANOVA and Turkey’s test for multiple comparisons.

**Supplementary figure 3.**
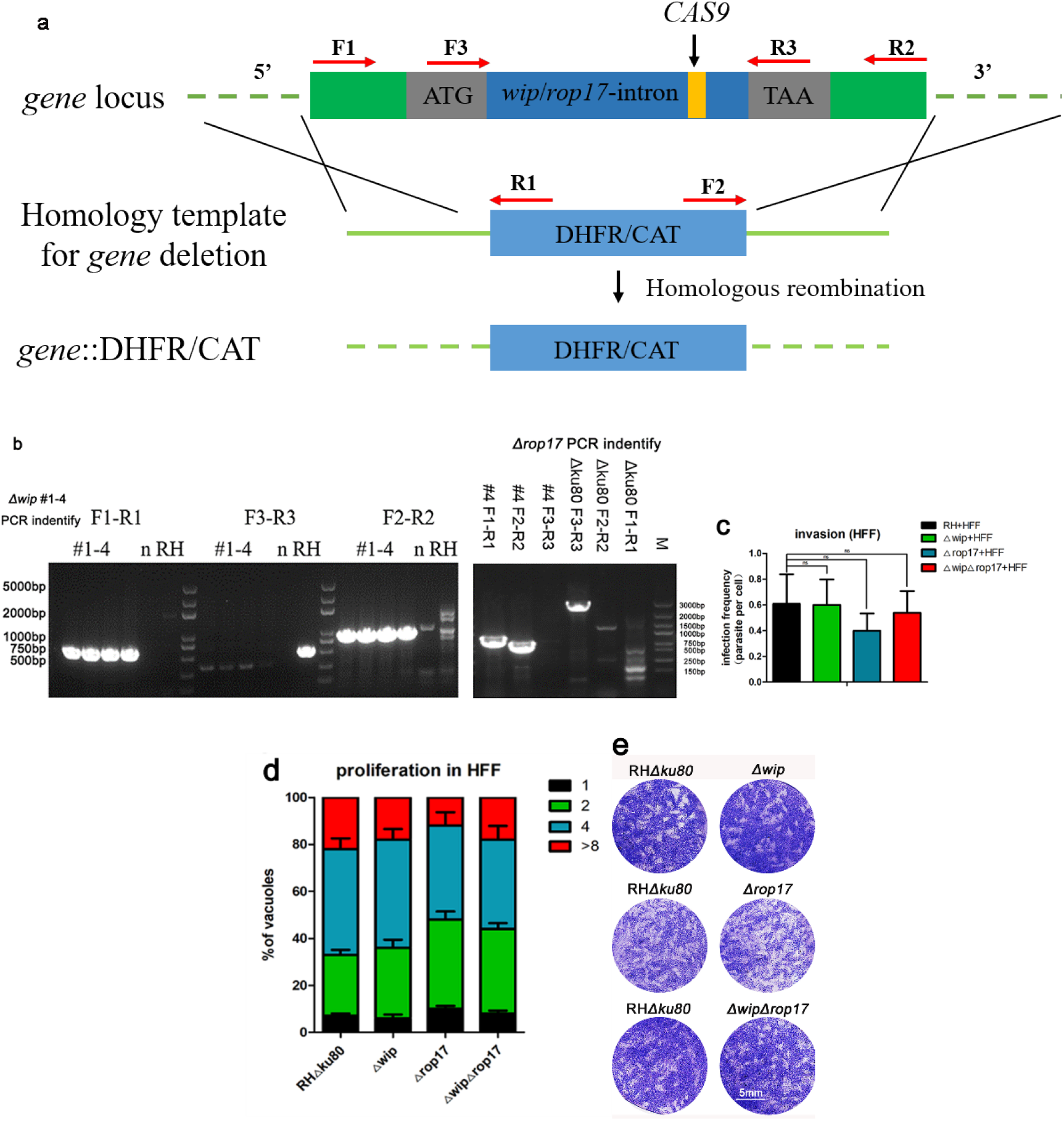
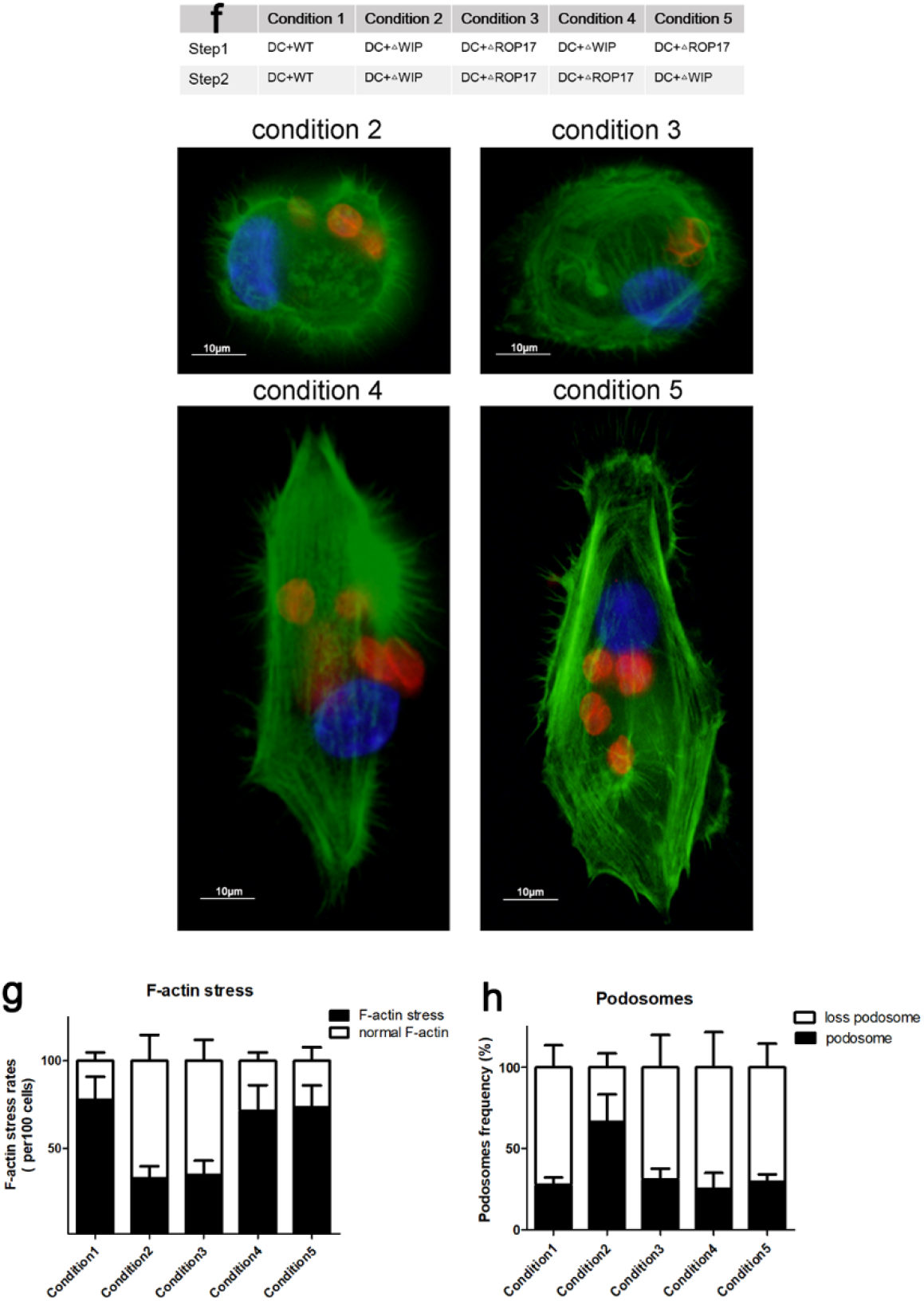
Determination of the phenotype of Tg*Δwip*, Tg*Δrop17* and Tg*ΔwipΔrop17* in vitro. (a) The schematic illustration of gene (*Δwip*, *Δrop17*) deletion. The guide RNA (yellow blot) was designed in the EuPaGDT library of TxoxDB. Target gene was replaced of *DHFR/CAT* gene, primers F1/R1and F2/R2 were designed to verify the integration. Primers F3/R3 were designed to identify the deletion of the target gene (b). (c) Invasion assay shows the same ability of invading HFF between parental and gene deletion strains. (d) Replication assay shows no significant differences proliferation between parental and gene deletion strains. (e) Plaque formation assay was used for detect the virulence of gene deletion strains. HFF were stained with crystal violet post inoculation 150 freshly tachyzoites for 7 days. (f) The F-actin structure phenotype of *T.gondii*-infected DCs. First, DCs were infected with parasites describe in Step1 condition (condition1 to condition5, MOI 3). 30 min after Step1 infection, uninvaded parasites were washed by PBS and infected with Step2 condition (condition1 to condition5, MOI 3) parasites respectively and observed DC actin structure. F-actin stress were stained with 488 Alexa Flour Phalloidin, the parasites were stained with anti-TgGAP45, and the nucleus were stained with Hoechest. Scale bars, 10 μm. Quantification of the percentage of DCs appearing podosomes (g) and F-actin stress fibers (h) on different condition were counted with 100 independent DCs. For all analyses, asterisks (*/**/***) indicate significant difference (\**p*<0.05, ** *p*<0.01, *** *p*<0.001), ns (no significant difference). Normally distributed data were repeated measured with one-way ANOVA, Tukey’s test for multiple comparisons.

**Supplementary figure 4.**
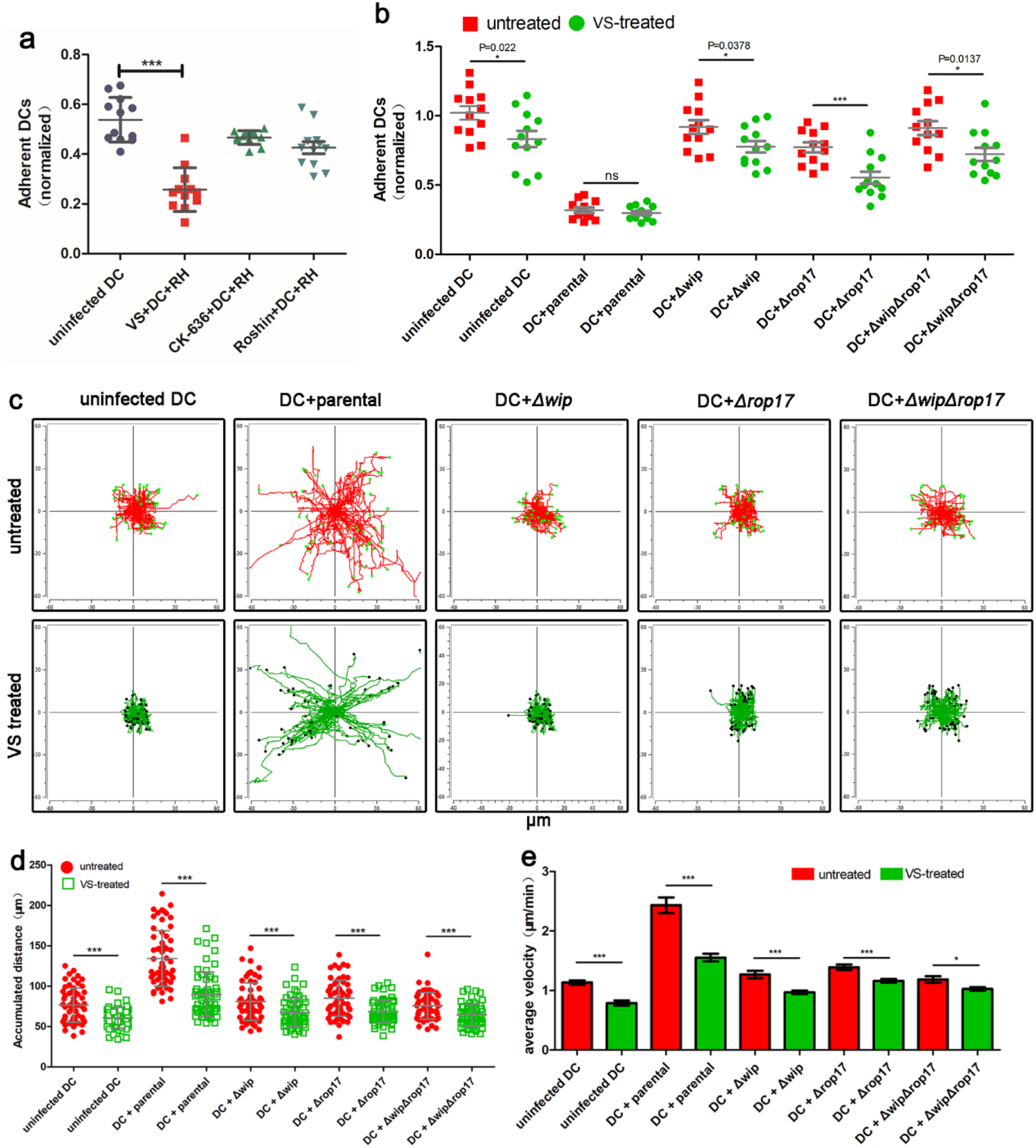
FAK inhibitor decrease DCs migration and adhesion on high stiffness substrate. (a) Cell adhesion of parental infected DCs treated with inhibitor VS-6063 (FAK inhibitor, 20μM), CK-636 (Arp2/3 inhibitor, 100μM) and Rhosin ( Rho A/B inhibitor, 100μM). *T.gondii* (parental, Tg*Δwip*, Tg*Δrop17 or* Tg*ΔwipΔrop17* parasites)-infected DCs treated with VS-6063 (FAK-inhibitor), and cell adhesion(b), motility plots (c), accumulated migrated distances (d) and cell average velocity (e) on high stiffness substrate were analyzed with one-way ANOVA, Dunnett’s multiple comparisons test.

**Supplementary Figure 5.**
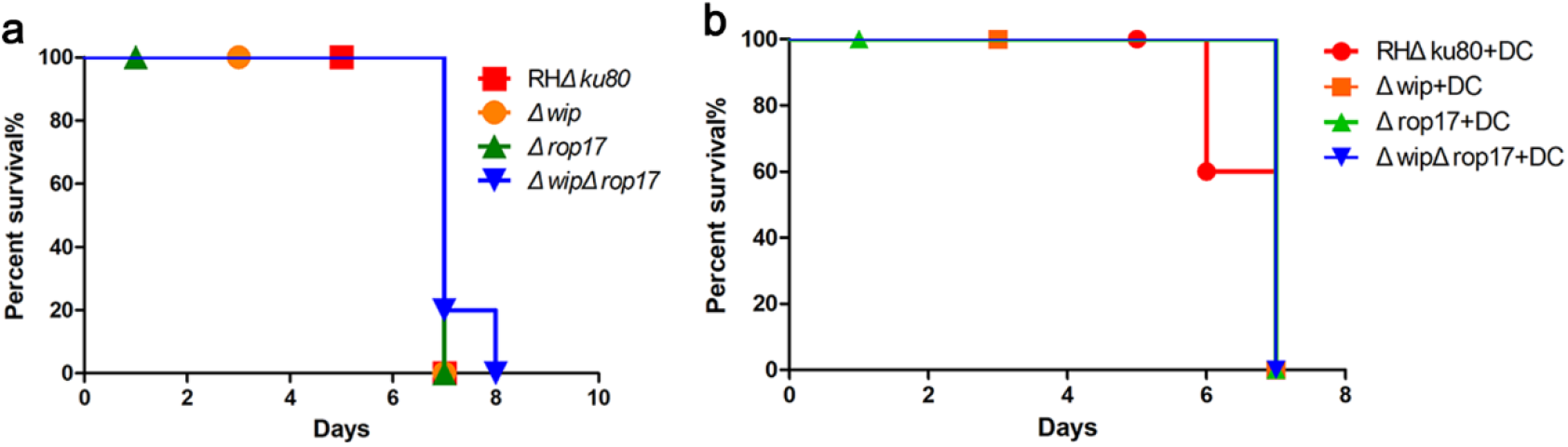
The virulence of gene deletion strains in mice. Growth curve of BALB/c mice i.p with 200 dose of freshly tachyzoites (a) or T.gondii infected-DCs (b) Non-significant difference were detected (Kruskal-Wallis test, P≥0.05).

**Table 1:**
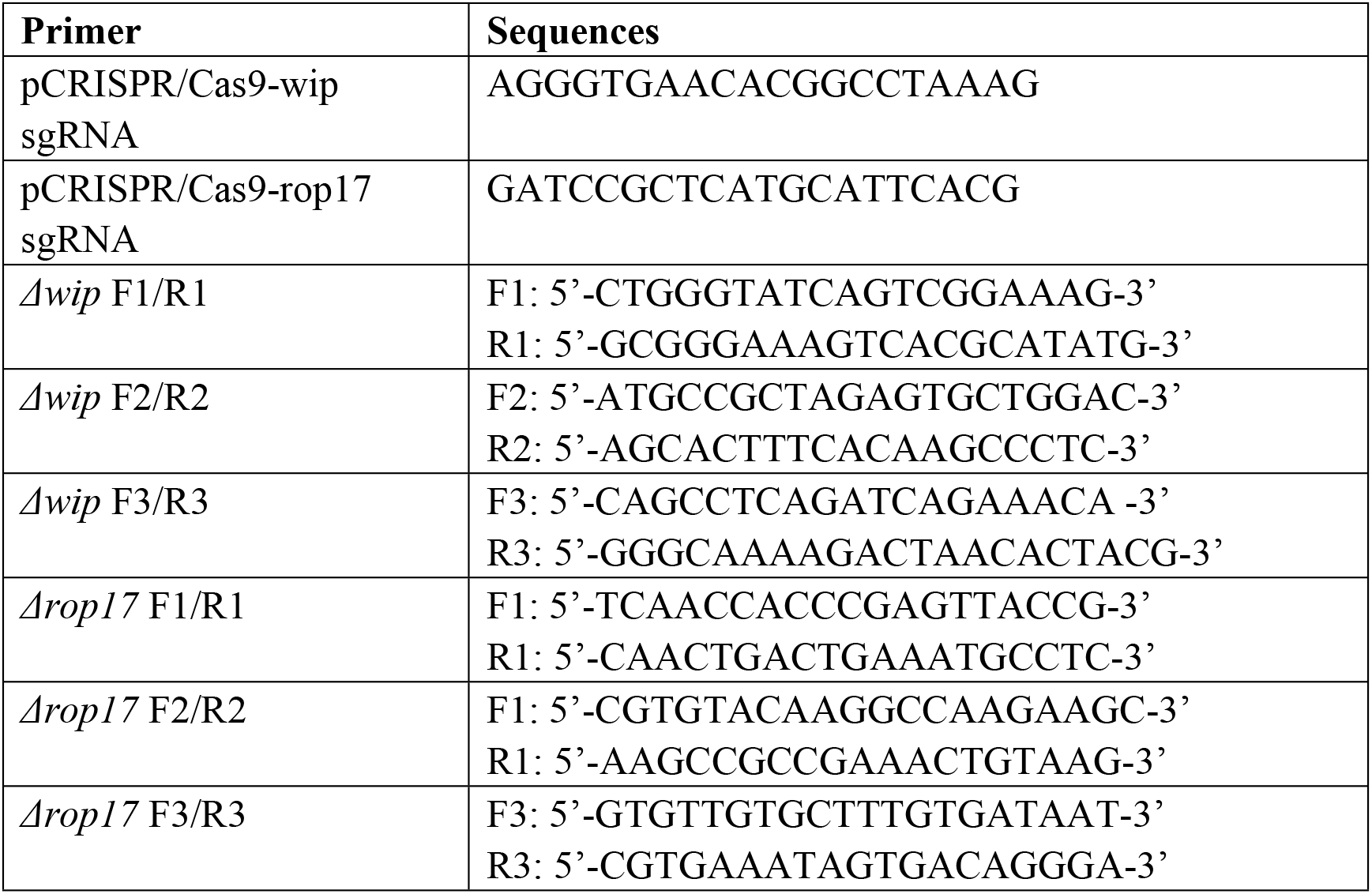
The sequences of gRNA and primers.

Key Resource Table.

**Key Resource Table**
| REAGENT or RESOURCE | SOURCE | IDENTIFIER |
| --- | --- | --- |
| **Antibody** |  |  |
| TgSAG1 (rabbit polyclonal) | Zhang H et al.,[50] | N/A |
| TgGAP45 (rabbit polyclonal) | Zhang H et al.,[50] | N/A |
| TgIMC1 (mouse polyclonal) | Li S et al.,[52] | N/A |
| FITC goat anti-mouse IgG(H+L) | Invitrogen | Cat#31569 |
| Cy3 goat anti-rabbit IgG(H+L) | Invitrogen | Cat#A10520 |
| Cy3 goat anti-mouse IgG(H+L) | Invitrogen | Cat#A10521 |
| Anti-mouse CD11a-FITC | Abcam | Cat#ab269336 |
| Anti-mouse CD11b-FITC | Abcam | Cat#ab269333 |
| Anti-mouse CD11c-FITC | Abcam | Cat#ab22540 |
| Anti-FAK (rabbit polyclonal) | Cell Signaling TECHNOLOGY | Cat#3283 |
| Anti-phospho-FAK (T397 )<br>(rabbit polyclonal) | Cell Signaling TECHNOLOGY | Cat#3285 |
| Anti-β actin (rabbit polyclonal) | Cell Signaling TECHNOLOGY | Cat#8457 |
| Anti-rabbit IgG-HRP (goat polyclonal) | Cell Signaling TECHNOLOGY | Cat#7074 |
| Anti-ZO-1 (mouse monoclonal) | Invitrogen | Cat#33-9100 |
| **Chemicals, peptides, and recombinant proteins** |  |  |
| Alexa Fluor™ 488 Phalloidin | Invitrogen | Cat#A12379 |
| Murine recombinant IL4 | PEPROTECH | Cat#214-14 |
| Murine recombinant GM-CSF | PEPROTECH | Cat#315-03 |
| Acrylamide | Sigma-Aldrich | Cat#A8887 |
| Bis-acrylamide | Sigma-Aldrich | Cat#146072 |
| Sulfo-SANPAH | Sigma-Aldrich | Cat#803332 |
| 3-aminopropyltrimethoxysilane | Sigma-Aldrich | Cat#A3648 |
| Poly-L-lysine | Sigma-Aldrich | Cat#P6282 |
| Hoechst | Sigma-Aldrich | Cat#23491-45-4 |
| VS-6063 (inhibitor) | MedChemExpress | Cat#HY-12289 |
| Rhosin (inhibitor) | MedChemExpress | Cat#HY-12646 |
| CK-636 (inhibitor) | MedChemExpress | Cat#HY-15892 |
| Cell Counting Kit-8 (CCK-8) | MedChemExpress | Cat#HY-K0301 |
| Fetal bovine serum (FBS) | Gibco | Cat#10100147 |
| DMEM | Gibco | Cat#11960044 |
| Phosphate-Buffered Saline (PBS) | Sigma-Aldrich | Cat#P4417 |
| Collagen type I | Sigma-Aldrich | Cat#C7661 |
| Collagenase type I | Gibco | Cat#17100017 |
| **Experimental models** |  |  |
| Mouse: C57BL/6 | Vital River Laboratories | 027 |
| Mouse: BalB/c | Vital River Laboratories | 028 |
| Mouse: CD-1 | Vital River Laboratories | 022 |
| Cell line: HFF | ATCC | SCRC-1041 |
| Cell line: Vero | ATCC | IM-C015 |
| *T.gondii* (RH) | Zhang H et al.,[50] | N/A |
| *T.gondii* (Pru) | Li S et al.,[52] | N/A |
| *T.gondii* (RH-mCherry) | Zhang H et al.,<sup>48</sup> | N/A |

